# Genotoxic chemotherapy impedes complement dependent cytotoxicity via Chk1-mediated CD59 regulation

**DOI:** 10.1101/2025.02.17.638751

**Authors:** Allison S.Y. Chan, Akshaya Anbuselvan, Patrick W. Jaynes, Charmaine Z.Y. Ong, Michal M. Hoppe, Wai Khang Yong, Vartika Khanchandani, Jie Min Lee, Nurulhuda Mustafa, Irfan Azaman, Phuong M. Hoang, Guo Hong, Wee-Joo Chng, Mark S. Cragg, Dennis Kappei, Claudio Tripodo, Anand D. Jeyasekharan

## Abstract

The DNA damage response (DDR) is a central regulator of cancer cell fate, coordinating both pro-death and pro-survival pathways in response to genotoxic stress. Here, we reveal an unexpected role for the DDR at the cell surface, in mediating immune evasion from complement-dependent cytotoxicity (CDC), an innate immune mechanism exploited by therapeutic monoclonal antibodies (mAbs). In the context of diffuse large B-cell lymphoma (DLBCL), where the anti-CD20 mAb rituximab utilizes CDC, we show that genotoxic chemotherapy induces expression of membrane-bound complement regulatory proteins (mCRPs) CD46, CD55, and CD59, thereby reducing CDC sensitivity and compromising rituximab activity. In this setting, CD59 emerged as the dominant DDR-induced inhibitor of complement-mediated killing. A high-throughput kinase inhibitor screen identified checkpoint kinase 1 (Chk1) as a critical mediator of this response. Mechanistically, DNA damage activates Chk1, enhancing CD59 transcription via an Sp1-bound promoter. Co-immunoprecipitation mass spectrometry revealed a Chk1 dependent remodelling of Sp1-associated complexes to a transcriptionally active state with recruitment of the histone acetyltransferase KAT2A. These findings expand the role of the DDR in immune resistance at the tumor cell surface, and highlight a negative interaction between chemotherapy and monoclonal antibodies that may require sequential administration or targeting of the Chk1– Sp1–CD59 axis.

**Significance:** The DNA Damage Response upregulates complement-protective proteins, extending its role in modulating immune evasion at the cell surface, with direct implications for combinations of chemotherapy and monoclonal antibodies widely used in cancer.

## INTRODUCTION

Genotoxic stress initiates the DNA damage response (DDR), a multifaceted signaling network that preserves genomic stability through DNA repair and regulated cell death. Beyond these canonical roles, emerging evidence underscores a critical, yet underexplored, function of DDR in modulating cell-extrinsic immunogenicity (1,2). DDR-driven changes in surface protein expression—such as upregulation of MHC-I, NKG2D ligands, and ICAM-1—can potentiate tumor cell recognition by cytotoxic lymphocytes, suggesting a dual role for DDR in both intrinsic genome preservation and extrinsic immune signaling (3–5).

However, the extracellular dimension of DDR remains incompletely characterized, especially in the context of post-chemotherapy tumor microenvironment dynamics and responsiveness to immunotherapy. To systematically identify DDR-regulated surface proteins, we previously employed a phage display-based ICOS screen, which revealed candidate antigens enriched on irradiated cells (6,7). Among these was Ly6D (6), and here we focus on another prominent DDR-induced target—a membrane-bound complement regulatory protein (mCRP)—to elucidate its immunological relevance and potential as a therapeutic modulator.

Complement is a component of innate immunity, functioning in immune surveillance through the promotion of cell lysis, chemotaxis, and opsonization upon activation (8). To prevent self-targeted complement activation, host cells express membrane-bound complement regulatory proteins (mCRPs), including CD46 (membrane cofactor protein), CD55 (decay-accelerating factor), and CD59 (protectin). CD46 and CD55 inhibit the formation and activity of C3/C5 convertases, thereby limiting complement cascade amplification (9,10) whereas CD59 inhibits terminal pathway activation by preventing the formation of the membrane attack complex (MAC), thereby protecting cells from lytic damage (11). Elevated expression of mCRPs has been observed across multiple malignancies, consistent with the hypothesis that tumors co-opt complement regulatory mechanisms to evade immune-mediated lysis (12).

Beyond their classical role in complement regulation, mCRPs also function as signaling receptors and immunomodulators. In cancer contexts, mCRPs have been shown to activate intracellular pathways, including Src and MAPK cascades (13,14), that drive tumor cell migration, invasion, and metastatic progression (15,16). Moreover, mCRPs interface with the adaptive immune response, influencing both tumor cell and T cell behavior in ways that promote immune evasion and tumor persistence (17).

Complement regulation is especially relevant in the context of complement-dependent immunotherapies. Several monoclonal antibodies (mAbs), including rituximab, elicit complement-dependent cytotoxicity (CDC) upon antigen binding, thereby leveraging host complement pathways to kill target cells. Rituximab, which targets CD20 on B-cell malignancies such as diffuse large B-cell lymphoma (DLBCL), is classified as a type I anti-CD20 antibody with strong CDC-inducing capacity (18,19). However, resistance to CDC has been linked to tumor-intrinsic upregulation of mCRPs, particularly CD55 and CD59, which inhibit complement activation and MAC formation. In preclinical and clinical settings, blockade of these mCRPs has been shown to enhance CDC and improve therapeutic efficacy (20,21). These observations raise the possibility that genotoxic treatments—by modulating mCRP expression—may inadvertently dampen the effectiveness of complement-fixing mAbs like rituximab.

Here, we identify a previously unrecognized mechanism by which DNA-damaging chemotherapy induces the transcriptional upregulation of mCRPs, thereby impairing mAb-mediated CDC and reducing therapeutic efficacy. To counteract this immunosuppressive adaptation, we conducted a targeted chemical screen and identified checkpoint kinase 1 (Chk1) inhibitors as potent suppressors of chemotherapy-induced mCRP expression. Notably, Chk1 inhibition restored rituximab-induced CDC activity in chemotherapy-treated tumor cells. Mechanistically, we demonstrate that Chk1 modulates the interaction between Sp1, a key transcription factor of CD59 transcription (22), and KAT2A, a histone acetyltransferase, in a DNA damage–dependent manner. These findings uncover a novel axis of therapeutic resistance involving Chk1-driven transcriptional control of complement regulators and suggest a strategy to enhance the synergy between genotoxic chemotherapy and immunotherapy through rational DDR pathway modulation.

## RESULTS

### mCRPs CD46, CD55, and CD59 are upregulated upon chemotherapy treatment

To identify cell surface antigens upregulated post-DNA damage, we conducted a phage-display antibody (Ab) screen comparing X-ray irradiated cells and non-irradiated cells (6) (**Fig. 1A Fig. S1A-C**). Using a published list of single-chain variable fragments (scFvs) that bound tumor-associated antigens as a frame of reference (23), several scFv sequences from our screen matched those previously identified as binding to membrane cofactor protein (MCP), also known as CD46. In a confirmatory experiment, X-ray irradiation of MCF10A cells displayed a dose-dependent increase of CD46 (**Fig. 1B**).

**Fig. 1.**
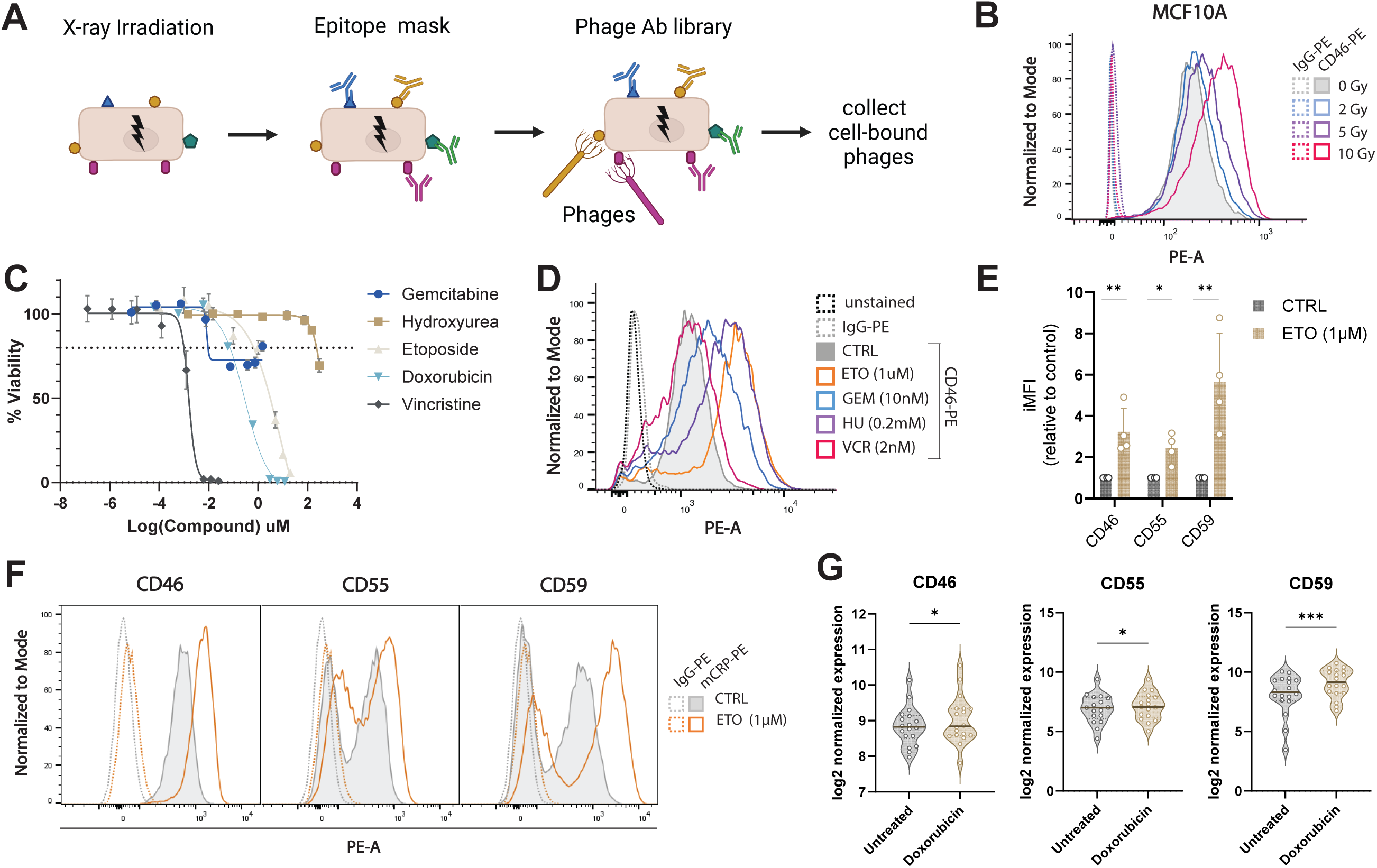
ICOS Screen identifies upregulation of CD46 as response to DNA damage **(A)** Schematic illustration of the multi-stage process of the phage-display antibody screen used to identify antibodies that preferentially bind to DNA-damaged cells. **(B)** Cell viability of SUDHL4 cells treated with indicated doses of compounds for 48 hours in technical triplicate. **(C)** Flow cytometry histogram of CD46 levels in MCF10A cells exposed to various doses of irradiation (0, 2, 5, 10 Gy) and stained with PE-conjugated anti-CD46 antibody or IgG-PE as isotype control. **(D)** Flow cytometry histogram of CD46 levels in SUDHL4 cells treated with various chemotherapy for 48H. Conditions include untreated control, IgG-PE isotype control, and treatments with 1uM etoposide (ETO), 10nM gemcitabine (GEM), 0.2mM Hydroxyurea (HU), and 2nM vincristine (VCR). **(E)** Representative flow cytometry histograms of CD46, CD55, and CD59 in SUDHL4 cells treated with or without etoposide (ETO). Cells were stained with PE-conjugated specific antibodies or IgG-PE as isotype control. **(F)** Quantification of relative integrated mean fluorescence intensities (iMFI) for CD46, CD55, and CD59 in SUDHL4 cells treated with ETO, normalized to untreated control. Statistical analysis was performed using ratio paired t-tests of IgG-corrected iMFI values. Data are presented as mean ± SD from three independent experiments. **(G)** Violin plot showing log2-normalized expression of CD46, CD55, and CD59 in 18 DLBCL cell lines (DB, Karpas422, RCK8, SUDHL10, U2932, WSUDLCL2, OCILY1, OCILY18, RI1, SUDHL2, SUDHL4, TOLEDO, CARNAVAL, DOHH2, NUDHL1, OCILY7, SUDHL5, SUDHL6) before and after adriamycin treatment. Solid lines indicate medians and dashed lines indicate quartiles. Statistical significance was determined using Wilcoxon tests. Statistical significance is denoted as follows: * p < 0.05, ** p < 0.01, *** p < 0.001.

Prompted by this observation, we broadened our analysis to other mCRPs. Functionally, mCRPs such as CD46, CD55, and CD59 protect host cells from complement-mediated lysis and are known to interfere with antibody-based cancer therapies by limiting complement-dependent cytotoxicity (CDC) (22). To examine this further, we employed a panel of diffuse large B-cell lymphoma (DLBCL) cell lines as a model for studying the effects of anti-CD20 monoclonal antibody rituximab (RTX) and often used in combination with chemotherapy (24). We observed that CD46 expression was significantly upregulated in the SUDHL4 DLBCL cell line after 48-hour treatment with genotoxic agents that induce either replication stress (gemcitabine, hydroxyurea) or direct DNA damage (etoposide, doxorubicin) (**Fig. 1C-D**), using doses of chemotherapy that evoked non-lethal DNA damage (80% viability). The microtubule-targeting agent vincristine did not upregulate CD46, supporting a direct link between DNA damage induction and CD46 expression.

Etoposide was selected as a representative genotoxic agent for subsequent studies due to its robust induction of CD46 and compatibility with flow cytometry. Flow cytometric analysis of SUDHL4 cells treated with etoposide revealed significant upregulation of CD46, CD55, and CD59, with CD46 and CD59 showing the strongest induction (**Fig. 1E-F**). To determine whether this effect extended to the transcriptional level, we analyzed a panel of 18 DLBCL cell lines treated with doxorubicin. Gene expression profiling (GSE224867) showed broad upregulation of mCRP transcripts, with CD59 exhibiting the most consistent and pronounced increase (**Fig. 1G**) (25).

### Chemotherapy impedes rituximab-induced CDC via upregulation of CD59

We hypothesized that etoposide-induced upregulation of mCRPs could impede the efficacy of rituximab, particularly through CD55 or CD59 (19,25). CD46 has shown a less pronounced CDC-enhancing effect compared to CD55 and CD59, possibly due to the insufficiency of complement regulation domain blockage by commercial anti-CD46 mAbs. Suitable cell line candidates for RTX-induced CDC were first identified using an in-vitro CDC assay, of which complement proteins are supplemented by normal human serum, and conversely where direct death can be ascertained with heat-inactivated serum (**Fig. 2A**). After 2 hours of incubation with normal human serum, cell viability after RTX treatment is then measured by Cell Titer Glo (CTG). RTX induced effective complement-mediated cell death across most DLBCL cell lines (SUDHL6, SUDHL4, SUDHL2, U2932, Karpas231, DOHH2, OCI-LY19, Karpas422) and not via direct cytotoxicity, consistent with previous reports (19,26) (**Fig. 2B**). Cell lines that had low levels of CD20 expression were insensitive to CDC (SC-1, HT) (**Fig. 2C**). We selected a CDC-sensitive cell line, SUDHL4, to test the effect of etoposide pre-treatment on RTX efficacy. In line with our hypothesis, RTX-induced CDC was impeded in the etoposide pre-treatment condition (ETO + RTX + human serum (HS)) compared to the control (RTX + HS) across the dynamic range (**Fig. 2D**). Direct cell death was not activated nor altered (RTX + heat inactivated HS (hiHS); ETO + RTX + hiHS). Using a RTX dose of 37 ng/mL, within the assay’s upper dynamic range, we confirmed that all genotoxic agents tested (gemcitabine, hydroxyurea, etoposide, and doxorubicin) significantly impeded CDC, whereas vincristine did not affect CDC levels (**Fig. 2E**).

**Fig. 2.**
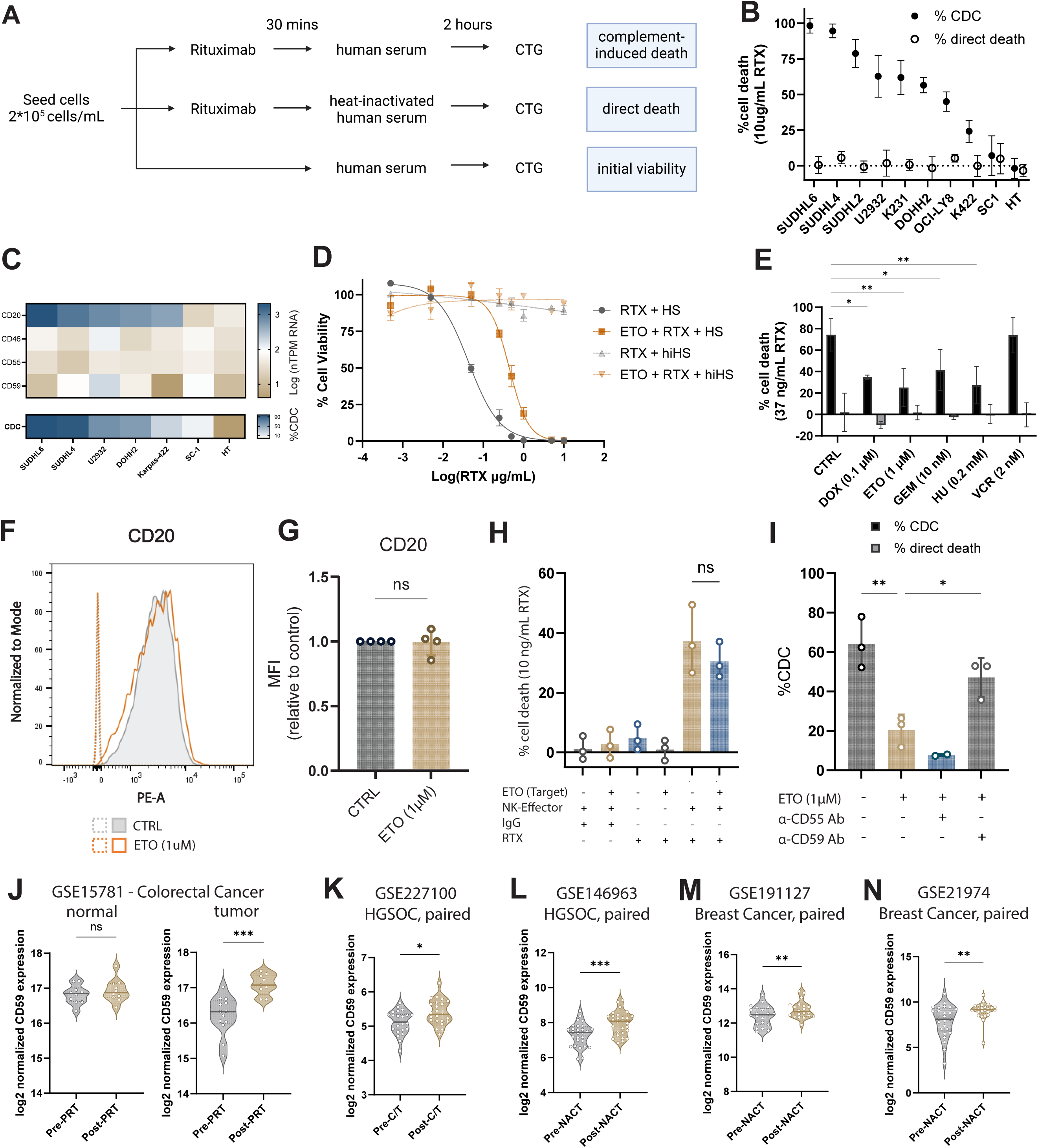
Chemotherapy prevents rituximab-induced CDC via upregulation of CD59 (**A**) Schematic illustrating the complement-dependent cytotoxicity (CDC) assay protocol. CTG, CellTiter-Glo. (**B**) Mean cell viability of 10 DLBCL cell lines treated with 10 ug/mL rituximab (RTX) over 2 hours, measured by CTG assay. Black dots represent CDC-mediated cell death, while clear dots indicate RTX-induced direct cell death. Error bars represent SD (n≥2 biological replicates). (**C**) Heatmap showing log-transformed normalized transcript per million (nTPM) RNA expression values for CD20, CD46, CD55, and CD59 in DLBCL cell lines, obtained from The Human Protein Atlas, ranked by %CDC activity derived from Fig. 2B. (**D**) Dose-response curve of RTX-induced cell death in SUDHL4 cells pre-treated with etoposide (ETO) for 48H, measured by CTG assay. HS, human serum; hiHS, heat-inactivated human serum. Data fitted using a four-parameter variable slope model (n=3 biological replicates). (**E**) Cell death induced by RTX (37 ng/mL) in SUDHL4 cells pre-treated with various chemotherapeutic agents: doxorubicin (DOX), etoposide (ETO), gemcitabine (GEM), hydroxyurea (HU), and vincristine (VCR) for 48H. Both CDC and direct cell death are shown (n=3 biological replicates). Statistical significance was calculated by 2-way ANOVA relative to respective control followed by uncorrected Fisher’s LSD. Significance of % CDC and not % apoptosis is shown. (**F**) Representative flow cytometry histogram of CD20 expression in SUDHL4 cells treated with or without ETO. Cells were stained with APC-conjugated anti-CD20 antibody or IgG-APC as isotype control. (**G**) Relative mean fluorescence intensity (MFI) of CD20 in SUDHL4 cells treated with ETO, normalized to untreated control (n=4 biological replicates). Statistical significance was calculated by ratio paired t-test of IgG-corrected MFI values. (**H**) Percentage of antibody-dependent cellular cytotoxicity in SUDHL4 cells. SUDHL4 target cells were treated with 1 µM etoposide for 48 hours prior to co-culture. Rituximab (10 ng/mL) or isotype control IgG was then added in the presence of NK92.CD16 effector cells to assess antibody-dependent cellular cytotoxicity (ADCC). Data are shown as mean ± SD of three biological replicates. (**I**) Percentage of CDC in SUDHL4 cells treated with 37 ng/mL RTX including ETO-treated with blocking antibodies against CD55, or CD59 (n=3 biological replicates). Statistical significance among groups was determined using a mixed effect model, followed by post hoc Dunnett’s multiple comparisons test. Data are presented as mean ± SD. Statistical significance is denoted as follows: * p < 0.05, ** p < 0.01, *** p < 0.001. (**J-N**) Violin plots showing CD59 expression in patient samples before and after treatment. (**J**) Unpaired tumor and normal tissue samples from colorectal cancer patients before and after preoperative radiochemotherapy (PRT) (GSE15781; unpaired t-test). (K) Paired HGSOC tumor samples before and after 6 cycles of carboplatin/paclitaxel (C/T) treatment (GSE227100). (L) Paired HGSOC tumor samples before and after neoadjuvant chemotherapy (NACT) (GSE146963). (M) Paired breast cancer samples before and after NACT (GSE191127). (N) Paired breast cancer samples before and after NACT (GSE21974). For paired analyses (K-N), statistical significance was determined using Wilcoxon tests. Solid lines indicate medians and dashed lines indicate quartiles (* p < 0.05, ** p < 0.01, *** p < 0.001).

As a control, we assessed the surface expression of the target antigen CD20 after etoposide treatment by flow cytometry, which showed no change (**Fig. 2F-G**). This confirmed that the observed chemo-upregulated mCRPs were a specific phenomenon correlated with reduced CDC in this context. Additionally, to determine whether the observed effects extended to other RTX-mediated cytotoxic mechanisms, we assessed antibody-dependent cellular cytotoxicity (ADCC), a complement-independent pathway. Using NK-92.CD16 cells as effectors, we observed a slight but non-significant reduction in cell death following etoposide treatment (**Fig. 2H**), suggesting that the antagonistic effect by chemotherapy treatment is likely restricted to modulation in complement activity. To test the role of CD55 and CD59 upregulation in Rituximab-induced CDC, these mCRPs were blocked with commercial neutralizing anti-CD55 (BRIC216) and anti-CD59 (YTH53.1) neutralizing antibodies, which were previously confirmed to block their targets at 10 µg/mL (27). Anti-CD55 (BRIC216) neutralization did not rescue CDC, demonstrating levels comparable to etoposide-treated cells (**Fig. 2I**). In contrast, the anti-CD59 (YTH53.1) neutralizing antibody significantly rescued CDC, suggesting that CD59 is the key mCRP for mediating CDC resistance upon chemotherapy treatment.

To evaluate the clinical relevance of CD59 upregulation, we analyzed *CD59* transcription levels in tumor samples from patients treated with neoadjuvant chemotherapy. Microarray analysis of colorectal cancer samples from patients receiving preoperative radiochemotherapy (PRT) revealed *CD59* upregulation specifically in tumor tissue, but not in matched normal tissue, indicating that this response is localized to the tumor and associated with treatment (GSE15781 (28), **Fig. 2J**). Additionally, analysis of paired tumor samples from patients with high-grade serous ovarian cancer (HGSOC) and breast cancer showed consistently elevated *CD59* expression following chemotherapy, compared to pre-treatment levels, across multiple independent datasets (GSE227100 (29), GSE146963 (30), GSE191127 (31), GSE21974 (32), **Fig 2K-N**). These findings support a consistent, treatment-induced upregulation of CD59 in clinical tumor samples, extending observations made in cell line models.

### Chk1 is a therapeutic target to recover rituximab-induced CDC efficacy

To investigate the mechanism underlying chemotherapy-induced CD59 upregulation and its negative impact on rituximab-mediated cytotoxicity, we conducted a targeted chemical screen to identify actionable pathways. Given that kinases are key regulators of cellular processes such as proliferation, metabolism, and signal transduction, we hypothesized that kinase signaling may mediate this adaptive response. To explore this, we utilized the Selleckchem kinase inhibitor library, which comprises 418 clinically tolerated and pharmacologically potent small-molecule inhibitors (**Fig. 3A**). Each compound was applied at 1 μM, with or without 1 μM etoposide co-treatment for 48 hours, prior to initiating the CDC assay (**Fig. 3B**). Compounds were then ranked based on normalized cell viability under etoposide-treated conditions relative to controls, allowing us to prioritize candidates that reversed chemotherapy-induced resistance to CDC (**Fig. 3C**). Ranking hits allowed us to select compounds that enhanced cell death with CDC, and those below the 5^th^ percentile of the screen (n=20 compounds) were mostly enriched in compounds targeting DNA damage response, such as Chk1 inhibitors and ATM/ATR inhibitors (**Fig. 3D**). Validation of the ATR inhibitor VE-821 and Chk1 inhibitors Rabusertib, MK-8776, and CCT244747 (SRA737) showed that Chk1 inhibitors specifically recovered CDC sensitivity when administered alongside etoposide treatment (**Fig. 3E**). CDC rescue by Rabusertib was apparent in SUDHL4 and in another CDC-sensitive cell line, SUDHL6 (**Fig. 3F 3G**). Notably, Rabusertib alone did not significantly alter CDC, suggesting Chk1’s specific involvement in the induced signaling mechanism.

**Fig. 3.**
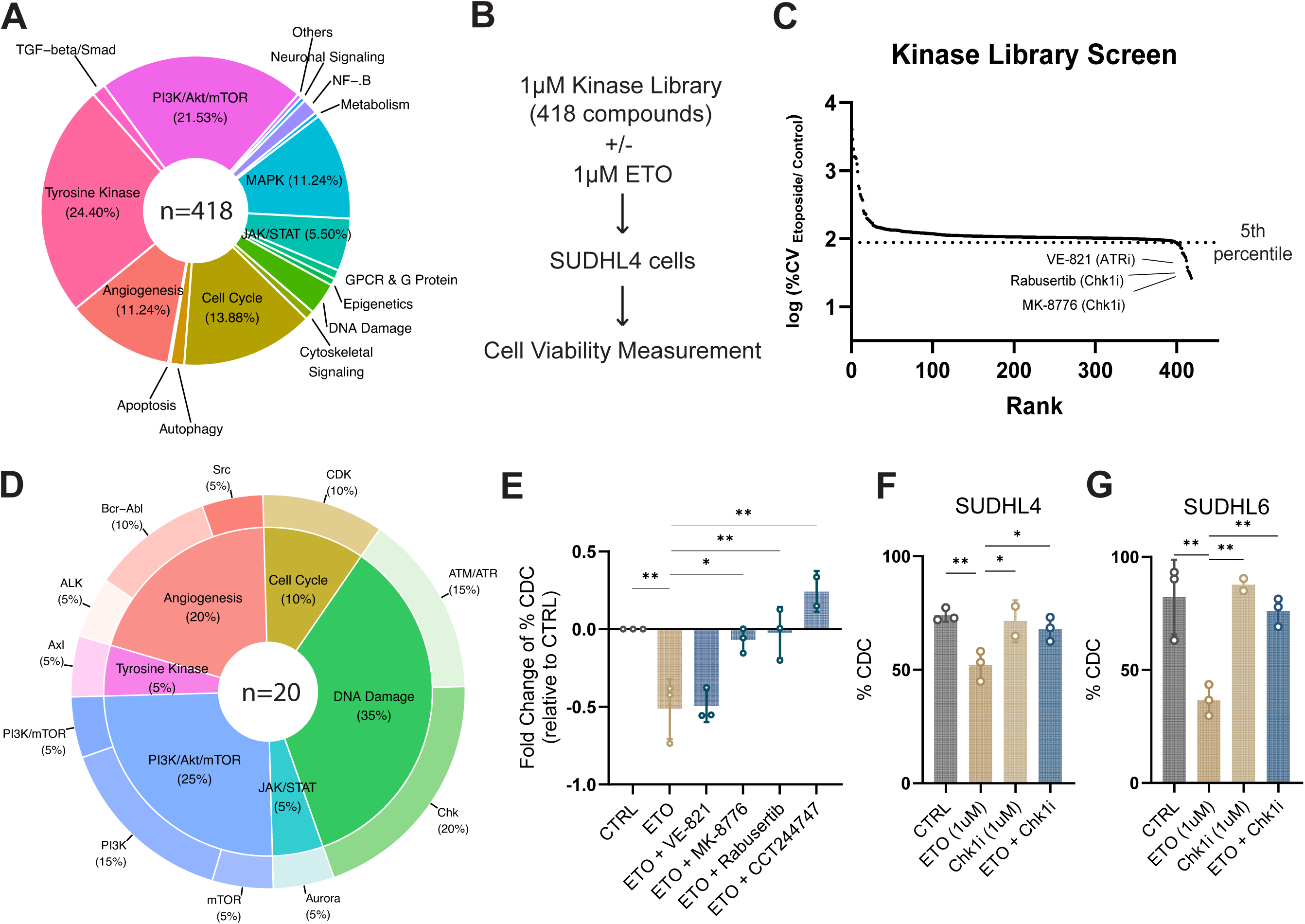
Kinase Library Drug Screen identifies Chk1 as therapeutic target to recover rituximab CDC efficacy **(A)** Schematic diagram illustrating the kinase library drug screen CDC procedure. **(B)** Pie chart depicting the distribution of targeted pathways in the 418 Selleckchem Kinase Library. **(C)** Ranked dot plot of kinase library screen hits, showing log % cell viability (CV) of etoposide-treated cells normalized to control. The dotted line indicates the 5th percentile threshold. **(D)** Nested pie chart illustrating the targeted pathways and corresponding protein targets among the hits below the 5th percentile. **(E)** Fold change of % CDC (normalized to control) in SUDHL4 cells treated with etoposide (ETO) and various ATR or Chk1 inhibitors (n=3 biological replicates). **(F, G)** % CDC in SUDHL4 (F) and SUDHL6 (G) cells pretreated with indicated compounds, including the Chk1 inhibitor Rabusertib, for 48 hours. For panels E-G, statistical significance was calculated by mixed-effects model with Dunnett’s multiple comparisons test relative to the ETO condition. Error bars represent SD of biological replicates. Statistical significance is denoted as follows: * p < 0.05, ** p < 0.01, *** p < 0.001.

Etoposide treatment activated Chk1, as indicated by the presence of phosphorylated Chk1 at Serine 345 (**Fig. 4A**). We thus hypothesized that activated Chk1 could be responsible for mediating the mCRP DNA damage response, and conversely, inhibiting Chk1 would block downstream signaling mediating this response. Indeed, combined treatment of etoposide and Rabusertib prevented the increase in mCRP levels in SUDHL4 cells (**Fig. 4B 4C**) and in other DLBCL cell lines (SUDHL6, SUDHL2, Karpas-231) (**Fig. 4D**). These results suggest that DNA damage induces Chk1-dependent regulation of mCRP levels.

**Fig. 4.**
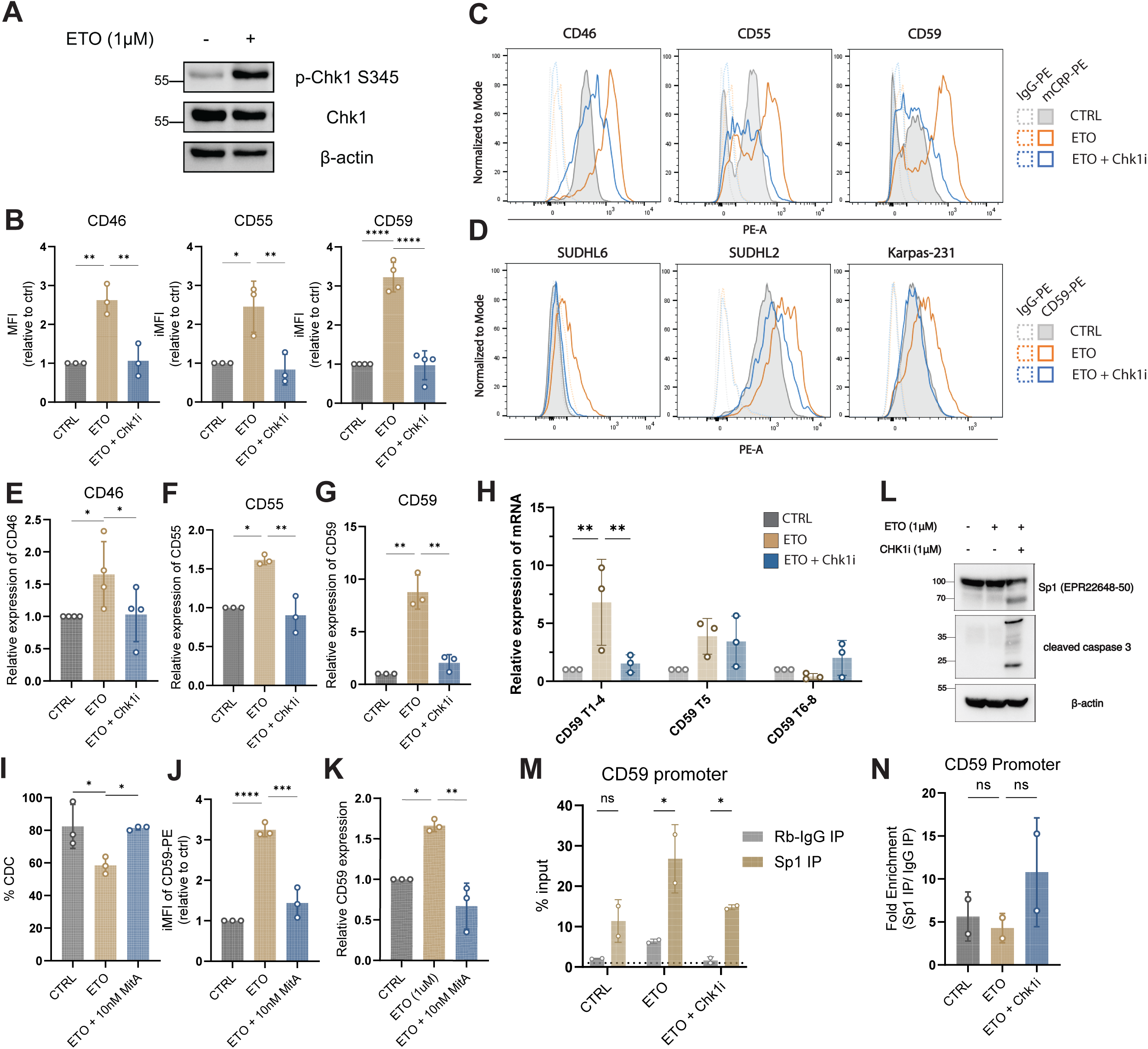
Chk1 is responsible for DDR-regulated mCRP expression **(A)** Immunoblot of Chk1 and phosphorylated Chk1 Ser345 (pChk1) in SUDHL4 cells treated with or without etoposide (ETO) for 48 hours. **(B)** Bar chart showing relative integrated mean fluorescence intensity (iMFI) of CD46, CD55, and CD59 in SUDHL4 cells treated with ETO alone or ETO + Chk1 inhibitor (Chk1i, Rabusertib). Statistical analysis was performed using repeated measures one-way ANOVA with Dunnett’s multiple comparisons test relative to the ETO condition. Error bars represent SD of 4 biological replicates. Statistical significance is denoted as follows: * p < 0.05, ** p < 0.01, *** p < 0.001. **(C)** Representative flow cytometry histograms in Fig. 4B of CD46, CD55, and CD59 expression in SUDHL4 cells under control conditions, ETO treatment, or ETO + Chk1i treatment. Cells were stained with PE-conjugated specific antibodies or IgG-PE as isotype control. **(D)** Flow cytometry histograms of CD59 expression in multiple DLBCL cell lines (SUDHL6, SUDHL2, Karpas-231) under control conditions, ETO treatment, or ETO + Chk1i treatment. Cells were stained with PE-conjugated anti-CD59 antibody or IgG-PE as isotype control. **(E-G)** Relative mRNA expression of *CD46* (E), *CD55* (F), and *CD59* (G) in SUDHL4 cells treated with etoposide (ETO) or ETO + Chk1 inhibitor (Chk1i, Rabusertib). **(H)** Relative mRNA expression of *CD59* exons T1-4, T5, and T6-8 in SUDHL4 cells after ETO or ETO + Chk1i treatment. **(I)** Percentage of complement-dependent cytotoxicity (%CDC) in SUDHL4 cells pretreated with ETO or ETO + 10 nM Mithramycin A (MitA). **(J,K)** Bar chart showing CD59 protein levels (MFI relative to control) (J) and relative *CD59* mRNA expression (K) in SUDHL4 cells treated with control, ETO, or ETO + 10 nM MitA. **(L)** Immunoblot analysis of Sp1 protein and cleaved caspase-3 in SUDHL4 cells after treatment with indicated compounds. **(M,N)** Chromatin immunoprecipitation (ChIP) in SUDHL4 cells, with DNA immunoprecipitation using an Sp1 antibody or control rabbit IgG. (M) Sp1 binding to the CD59 promoter is shown as % input. (N) Fold enrichment of Sp1 binding was calculated as the ratio of signal from Sp1 IP relative to IgG IP. For all panels, statistical significance was calculated by ANOVA followed by post hoc Dunnett’s multiple comparisons test relative to the ETO condition. Error bars represent SD of biological replicates. Statistical significance is denoted as follows: * p < 0.05, ** p < 0.01, *** p < 0.001.

### Chk1 regulates mCRP transcription via Sp1-dependent transcription

Examining this phenomenon at the transcriptional level, we observed that transcripts of *CD46*, *CD55*, and *CD59* were upregulated in SUDHL4 cells after etoposide treatment, and that this upregulation was prevented by the addition of Rabusertib (**Fig. 4E, 4F, 4G**). This suggests that Chk1 regulates mCRP expression at the transcriptional level. We next examined potential transcriptional regulators downstream of Chk1 signaling, referencing a previous study by Du et al. that identified key transcription factors responsible for *CD59* expression (**Fig. S2D**) (22). Nf-κB and Sp1 control expression of Exon 1 (T1-4), while TP53 drives Exon 1’ (T5) and CREB drives exon 1’’ (T6-8). Our results show that etoposide primarily upregulates *CD59* expression at Exon 1 (T1-4), with no significant change in Exon 1’ (T5) or Exon 1’’ (T6-8) expression (**Fig. 4H**). Rabusertib consistently reversed expression at Exon 1 (T1-4) in SUDHL4 and SUDHL6 cells, suggesting that NF-κB or Sp1 could be primarily responsible for the altered expression (**Fig. 4H Fig. S2B**).

It has been proposed that NF-κB primarily drives inducible expression of *CD59*, whereas Sp1 maintains constitutive expression (22). However, our experiments with NF-kB pathway inhibitors (QNZ, Dexamethasone, MLN120B, JSH-23) failed to rescue CDC following etoposide exposure (**Fig. S2C**). In contrast, treatment with Mithramycin A, a selective inhibitor of GC-rich DNA-binding transcription factors such as Sp1 successfully rescued CDC in both SUDHL4 and SUDHL6 cells (**Fig. 4I S2D**). This treatment also prevented the etoposide-induced upregulation of CD59 at both the protein and transcript levels (**Fig. 4J 4K**). Consistent with this, Sp1 binding sites are present in all three mCRP promoters (33), in line with supporting evidence that Mithramycin A prevented the induced *CD46* expression (**Fig. S2E**). These results suggest that the observed upregulation of mCRPs occurs through DNA damage-induced activity at the Sp1-bound site. Concordantly, treatment with Mithramycin A did not reduce basal mCRP expression, affirming that the targeted site primarily regulates the inducible expression of mCRPs (**Fig. S2F**).

Interestingly, this increase in mCRP expression is unlikely to be driven by elevated Sp1 protein levels, as Sp1 abundance remained unchanged after etoposide treatment (**Fig. 4L**). However, co-treatment with etoposide and the Chk1 inhibitor Rabusertib led to a decrease in full-length Sp1 and the appearance of a ∼70 kDa cleavage product, consistent with caspase-mediated cleavage of Sp1 (34) (**Fig. 4L**).

We next asked whether the upregulation of CD59 could be attributed to increased Sp1 recruitment to the CD59 promoter. To test this, we designed primers targeting a 248 bp region of the CD59 promoter and performed ChIP-qPCR using either anti-Sp1 or control IgG. Surprisingly, there was a modest increase in non-specific IgG binding following etoposide exposure (**Fig. 4M**), potentially reflecting increased chromatin accessibility or nonspecific protein-DNA interactions following DNA damage. While normalization to IgG control revealed no significant increase in Sp1 enrichment at the CD59 promoter (**Fig. 4N**), this normalization may obscure true biological changes in Sp1 occupancy, given that both specific and non-specific antibody access likely increase in a more permissive chromatin environment. These findings raise the possibility that Sp1 is still functionally active at the promoter, and that altered transcriptional output may result from changes in chromatin architecture or the recruitment of specific co-factors, rather than from increased promoter binding alone.

### DNA damage remodels the Sp1 complex in a Chk1-dependent manner to promote mCRP expression

Based on this, we hypothesized that DNA damage may instead remodel the Sp1-associated transcriptional complex. To investigate this, we performed label-free quantitative co-immunoprecipitation mass spectrometry (Co-IP LFQ-MS) using a Sp1-specific antibody (EPR22548) in SUDHL4 cells treated under three conditions: untreated (CTRL), etoposide (ETO), and etoposide with Chk1 inhibition (ETO+Chk1i), each in four biological replicates (**Fig. 5A**). Proteins significantly enriched in Sp1 pulldowns compared to IgG controls were identified using a threshold of mean log2 fold enrichment (FE) > 2 and p-value < 0.05. Fifteen proteins were consistently enriched across all three conditions, while six were uniquely enriched in the ETO condition (**Fig 5B**). Notably, the ETO+Chk1i interactome overlapped entirely with that of CTRL, suggesting that Chk1 activity is required for the altered Sp1-associated complex observed in response to DNA damage. Principal component analysis (PCA) further supported a difference in SP1 interactomes, where clear separation was observed between Sp1 and IgG pulldowns in both the CTRL and ETO+Chk1i conditions, whereas ETO-treated samples showed less separation (**Fig. 5C**), possibly indicating a weaker or more transient set of Sp1 interactions following DNA damage. Sp1 itself was the most highly enriched protein in all conditions (**Fig. 5D**) confirming the specificity of the pulldown.

**Fig. 5.**
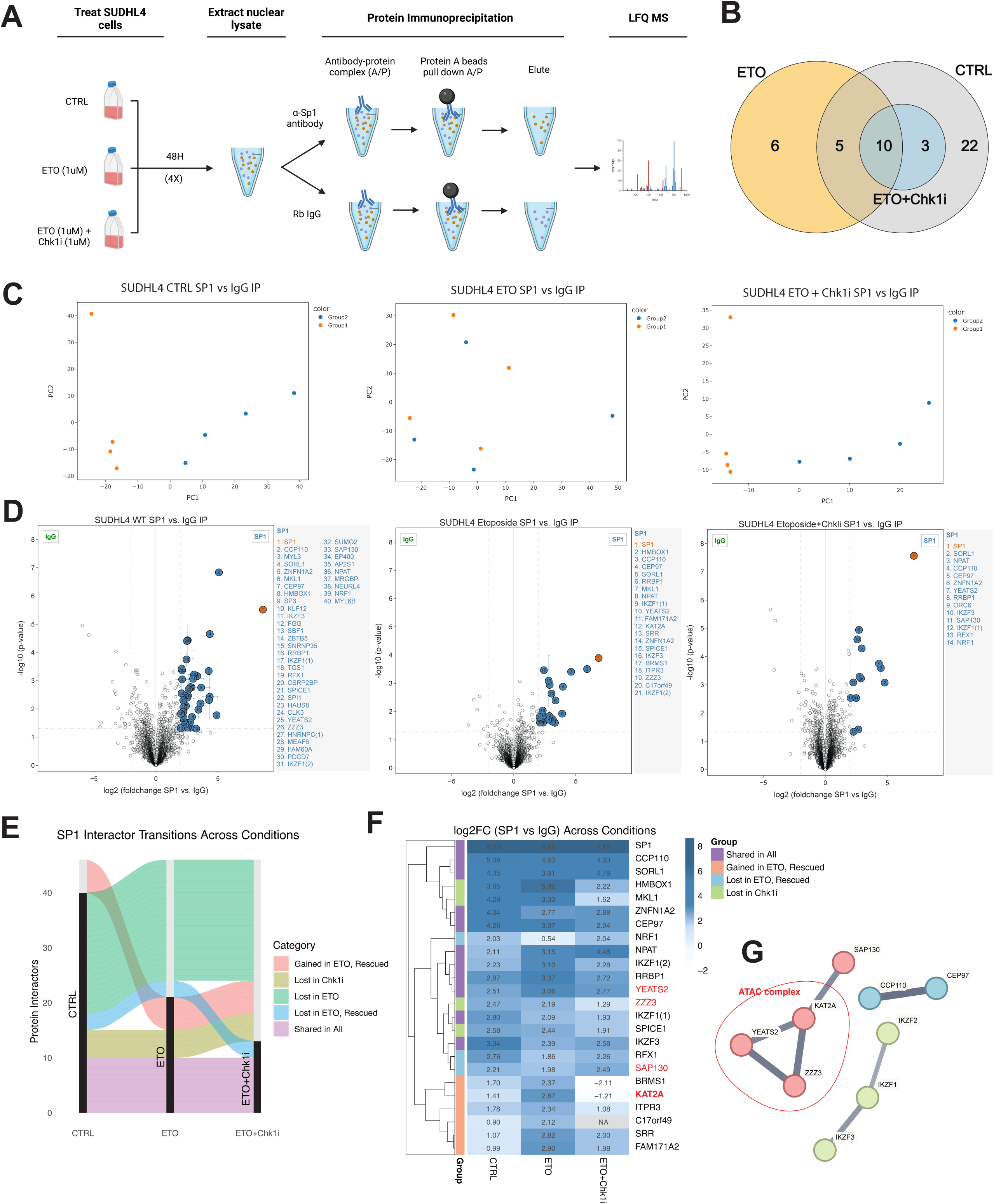
Etoposide treatment induces Chk1-dependent changes to the Sp1 interactome **(A)** Schematic diagram of co-immunoprecipitation mass spectrometry workflow. SUDHL4 cells were treated under three conditions (CTRL, ETO, ETO + Chk1i (Rabusertib)) for 48 hours in 4 biological replicates. Nuclear lysates were incubated with either Sp1 antibody (EPR22648) or rabbit IgG control. Protein A beads were used to pull down antibody-protein complexes, followed by protein elution and label-free quantification mass spectrometry (LFQ-MS) analysis. **(B)** Venn diagram showing significant Sp1-interacting proteins (vs IgG) under each condition (log2 fold change > 2 and p-value < 0.05). **(C)** Principal component analysis (PCA) plots of pulldown samples. Group 1: Sp1; Group 2: IgG. **(D)** Volcano plots showing proteins enriched in Sp1 IP compared to IgG control under CTRL, ETO, and ETO+Chk1i conditions. Significant hits were defined using a dual threshold of log2 fold change > 2 and p-value < 0.05. Error bars represent the standard deviations from three iterations of zero-value imputation. **(E)** Sankey diagram illustrating the overlap of Sp1-interacting proteins across conditions, categorized by interaction dynamics. **(F)** Heatmap showing log2 fold changes of selected Sp1-associated proteins grouped by interaction pattern: “Shared in All”, “Gained in ETO, Rescued”, “Lost in ETO, Rescued”, or “Lost in Chk1i”. **(G)** STRING protein-protein interaction network of selected proteins from panel (F). Edges represent physical interactions or functional associations with a minimum required confidence score of 0.6, derived from text mining, experimental data, and curated databases.

Proteins that were lost in ETO and not rescued by Chk1 inhibition were excluded from further analysis, as they likely represent interactions disrupted independently of DNA damage response (**Fig. 5E 5F**). Instead, we focused on interactions gained or lost in ETO and reversed by Chk1 inhibition, on a background of shared protein interactions. We therefore focused on the other significant proteins, of which STRING protein interaction network analysis of these selected proteins revealed several functionally relevant complexes (**Fig. 5G**). STRING protein-protein interaction network analysis of these candidates revealed several biologically relevant complexes (**Fig. 5G**). Notably, transcription factors IKZF1, IKZF2 (ZNF1A2), and IKZF3 were consistently associated with Sp1 across conditions, suggesting a potential transcription factor partnership. Components of the ATAC chromatin remodeling complex, including YEATS2 and ZZZ3, also showed stable interactions with Sp1. However, the histone acetyltransferase KAT2A, member of the KAT module within the ATAC complex, was significantly enriched only in the ETO condition. KAT2A catalyzes histone H3 lysine 9 acetylation (H3K9ac), a mark associated with transcriptional activation (35). In contrast, the interaction between Sp1 and SAP130, a component of the Sin3A-HDAC corepressor complex, was reduced following ETO treatment (36). This shift suggests a remodeling of the Sp1 complex to an activating transcriptional state in response to DNA damage.

Together, these findings support a model in which DNA damage leads to a reconfiguration of the Sp1 transcriptional complex, characterized by increased recruitment of transcriptional co-activator KAT2A and loss of repressor SAP130, thereby promoting mCRP gene expression. Inhibition of Chk1 reverses this remodeling, potentially restoring a repressive transcriptional environment and sensitizing cells to complement-dependent cytotoxicity (**Fig. 6**).

**Fig. 6.**
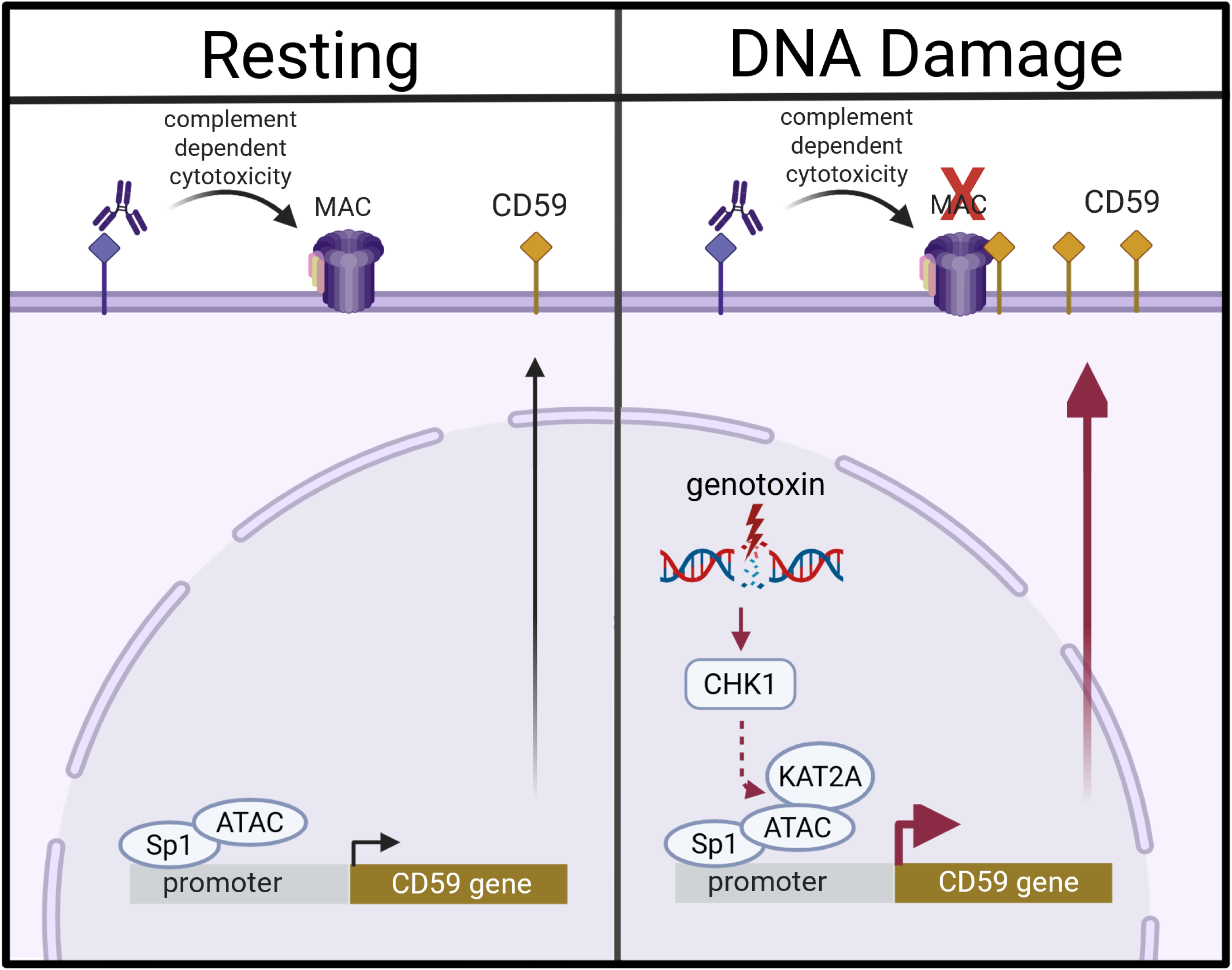
Illustration of chemotherapy-induced rituximab antagonism via CD59 upregulation. Schematic representation of the mechanism by which genotoxic agents antagonize rituximab-mediated complement-dependent cytotoxicity (CDC) through CD59 upregulation. Left panel: Under normal conditions, rituximab can induce CDC, resulting in membrane attack complex (MAC) formation and cell lysis. CD59 expression is driven by Sp1. Right panel: In the presence of DNA damage, genotoxic agents activate Chk1. Activated Chk1 associates KAT2A with ATAC complex and Sp1, driving increased mCRP gene expression, particularly CD59. Elevated CD59 levels block MAC formation, inhibiting rituximab-driven CDC.

## DISCUSSION

This study identifies a previously unrecognized facet of the DNA damage response that modulates expression of membrane complement regulatory proteins. While DDR is classically associated with intrinsic cellular processes such as cell cycle arrest, DNA repair, and apoptosis, our findings reveal that it also governs cell-extrinsic immune modulators, including CD46, CD55, and CD59, which function to inhibit complement activation. This adaptive response may help limit collateral tissue damage during stress but may also be co-opted by tumor cells to evade immune clearance. In the context of diffuse large B-cell lymphoma, we demonstrate that DNA-damaging agents like etoposide antagonize rituximab-induced complement-dependent cytotoxicity (CDC), primarily through induction of CD59 (27,37).

To define the regulatory pathway underlying mCRP induction, we performed a kinase inhibitor screen and identified Chk1 as a key modulator of mCRP expression following DNA damage. Pharmacological inhibition of Chk1 using Rabusertib reversed chemotherapy-induced CD59 expression and restored CDC activity. Mechanistically, we found that the mCRP response depends on the transcription factor Sp1, as the Sp1-inhibitory compound Mithramycin A suppressed DNA damage–induced CD59 and CD55 expression. However, etoposide treatment did not increase Sp1 protein levels or specific promoter occupancy, suggesting that transcriptional activation is driven not by enhanced DNA binding, but rather through altered cofactor interactions or post-translational modification. Using co-immunoprecipitation and mass spectrometry, we identified a DNA damage–induced interaction between Sp1 and the histone acetyltransferase KAT2A. This interaction likely facilitates the recruitment of the ATAC complex, promoting local chromatin acetylation and transcriptional activation of mCRP genes. Notably, this Sp1–KAT2A interaction was abrogated by Chk1 inhibition, indicating that Chk1 activity is required for the assembly or stability of this transcriptional complex.

Together, these data identify a novel Chk1 signaling axis that links the DNA damage response to mCRP expression through transcriptional regulation by KAT2A and Sp1. This insight provides a mechanistic rationale for targeting Chk1 to overcome immune resistance induced by genotoxic therapies.

The therapeutic relevance of this pathway is underscored by recent clinical interest in Chk1 inhibitors (38). Recently, CCT245737 (SRA737), for example, demonstrated good tolerability and a favorable pharmacokinetic profile in a Phase I/II clinical trial (NCT02797964) (39). While single-agent potency has been limited, combinations with chemotherapy agents have shown promise, such as low dose gemcitabine (NCT02797977) (40). Studies of Chk1 inhibitors with chemotherapy agents (e.g., etoposide, gemcitabine) have also confirmed strong synergistic anticancer effects in cancer cell line models (41,42). In our study, co-treatment with SRA737 restored rituximab-induced CDC after etoposide exposure, supporting a dual benefit: enhanced cytotoxic synergy and restored antibody-mediated clearance. In parallel, our findings suggest therapeutic value in neutralizing mCRPs directly—either with CD59/CD55-blocking antibodies or through engineered antibody formats (e.g., bispecifics, ADCs) that circumvent mCRP-mediated resistance .

Beyond direct complement-mediated lysis, DDR-induced mCRP upregulation may shape the broader immune landscape. For instance, CD55 blocks the generation of anaphylatoxins C3a and C5a, which, while classically pro-inflammatory, have been shown in many tumor models to recruit immunosuppressive populations such as MDSCs and M2 macrophages and to inhibit NK or cytotoxic T-cell infiltration (43). Thus, while mCRP-mediated suppression of complement may protect normal tissues, it could also foster an immunosuppressive tumor microenvironment. Supporting this, a recent study reported that CD55 suppresses the generation of ICOSL+ B cells following chemotherapy, thereby dampening anti-tumor immunity (44). Our observation that CD55 is upregulated following genotoxic stress suggests a second mechanism, beyond CDC inhibition, by which chemotherapy may impair adaptive anti-tumor responses, and highlights the therapeutic relevance of CD55 blockade (45).

While our in vitro findings offer compelling mechanistic insights, additional work is required to determine their translational relevance. Future studies should evaluate the kinetics and persistence of mCRP upregulation following chemotherapy in vivo. This will require longitudinal sampling of tumor tissue or liquid biopsies from patients receiving genotoxic agents to determine whether transient mCRP upregulation coincides with windows of reduced monoclonal antibody efficacy. Preclinical modeling of this phenomenon in mice is inherently challenging. Murine complement activity is roughly half that of human complement and mCRPs act in a species-selective manner, meaning that human mCRPs expressed on xenografted tumors may not effectively inhibit mouse complement (27,45). Moreover, mice possess a distinct complement regulatory landscape, including CD55 and CD59 homologs, and the rodent-specific Crry, which combines functions of human CD46 and CD55 and serves as their major C3 convertase regulator. Differences in the expression patterns and inhibitory potency of these molecules, along with reduced baseline complement activity, make it unlikely that standard mouse models fully recapitulate the combined effects of genotoxic chemotherapy, mCRP modulation, and monoclonal antibody activity observed in patients. Syngeneic or humanized mouse models, ideally incorporating homologous complement and mCRP systems, will therefore be essential for dissecting how DNA damage response–induced mCRP expression alters immune cell infiltration, complement activation, and antibody efficacy within an intact immune system.

In summary, our study identifies a novel Chk1–Sp1–CD59 axis through which DNA damage induces mCRP expression and confers resistance to complement-dependent cytotoxicity. These findings enhance the prevailing view of cell-intrinsic DNA damage responses and reveal a mechanism by which genotoxic therapies can modulate immune resistance. Our work provides strong rationale for improving therapies that combine genotoxins with immunotherapies through strategic combinations of Chk1 inhibition or CD59/CD55-targeted agents.

## MATERIALS AND METHODS

### Cell culture and reagents

MCF10A cells were cultivated in DMEM/F-12 (Dulbecco’s Modified Eagle Medium/Nutrient Mixture F-12) medium (Invitrogen) containing 5% horse serum, 10 μg/ml insulin, 20 ng/ml EGF, 100 ng/ml choleratoxin, 500 ng/ml hydrocortisone and 1% pen/strep and grown at 37°C and 5% CO_2_. Human DLBCL cell lines (DOHH2, HT, Karpas-231, Karpas-422, OCI-LY8, SC-1, SUDHL2, SUDHL4, SUDHL6, U2932) were cultured in RPMI-1640 (Roswell Park Memorial Institute 1640) medium supplemented with 20% heat-inactivated fetal bovine serum (FBS, HyClone Research Grade Fetal Bovine Serum, South American Origin) and grown at 37 °C and 5% CO_2_. MCF10A, DOHH2, HT, SUDHL4, and SUDHL6 cell lines were authenticated with STR profiling by ATCC Cell Authentication. Cells were confirmed mycoplasma-free by MycoAlert^®^ Mycoplasma Detection Kit (Lonza).

To maintain surface antigens prior to screening, MCF10A cells were harvested using enzyme-free cell dissociation buffer (Millipore), or StemPro Accutase Cell Dissociation Reagent (Invitrogen). Irradiation was performed at 3mA/130v on a Faxitron Cabinet X-Ray generator, with duration of exposure adjusted to define dosage. Compounds used are as follows: CCT244747 (MedChemExpress, HY-18175), CPI-537 (MedChemExpress, HY-100482), Curcumin (MedChemExpress, HY-N0005), Dexamethasone (MedChemExpress, HY-14648), Doxorubicin (MedChemExpress, HY-15142), Etoposide (MedChemExpress, HY-13629), Gemcitabine (MedChemExpress, HY-17026), Hydroxyurea (Sigma-Aldrich, H8627), JSH-23 (MedChemExpress, HY-13982), Mithramycin A (Sigma-Aldrich, M6891), MK-8775 (MedChemExpress, HY-15532), MLN120B (MedChemExpress, HY-15473), QNZ (MedChemExpress, HY-13812), Rabusertib (MedChemExpress, HY-13812), VE-821 (MedChemExpress, HY-14731), Vincristine (Adooq Bioscience, A10973). For all drug treatment experiments, cells were plated at a ratio of 10^6^ cells per 5mL media and treated with compounds for 48 hours. For treatment with Mithramycin A, cells were treated for 16 hours prior to collection due to cytotoxicity.

### Cell lysis and immunoblotting

Cells were lysed in RIPA buffer (Thermo Scientific) containing 0.2mM MgCl2, 0.01U/ µL BenzonaseⓇ Nuclease (Merck Millipore), 1X protease inhibitor (Sigma), 1X phosphatase inhibitor (Sigma). Protein concentrations were determined by BCA assay (Thermo Scientific) and equal concentrations were diluted into NuPage^TM^ LDS Sample Buffer and DTT. After electrophoresis, proteins were transblotted to PVDF membranes (Thermo Scientific). Blots were blocked with 5% skim milk/ TBS-Tween followed by incubation with primary antibodies overnight at 4°C. The blots were incubated with anti-rabbit IgG, HRP-linked antibody (Cell Signaling Technologies, #7074) and anti-mouse IgG, HRP-linked antibody (Cell Signaling Technologies, #7076) and were then developed using Immobilon HRP substrate (Merck Millipore). Blots were visualized using a ChemiDoc Imaging System (Bio-rad). Antibodies specific for the following proteins were used for immunoblotting: Cleaved caspase 3 (Asp 175) (Cell Signaling Technologies #9661), Chk1 (Cell Signaling Technologies, #2360), phospho-Chk1 Ser345 (Cell Signaling Technologies, #2348), β-Actin (Sigma-Aldrich, #A2228), Sp1 [EPR22648;50] CHIP grade (Abcam, ab231778).

### Chemical screening

Chemical screening was performed at the Center for High-throughput Phenomics, Genome Institute of Singapore. Compounds from Selleckchem Kinase Inhibitor Library (418 compounds) were printed at 1µM on a 384-well tissue culture plate (Corning) with an acoustic liquid handler (Echo, Beckman Coulter Life Sciences). Identification of accurate hits was ensured by assessing plate uniformity and Z’ score prior to procedure (46). Edge effect was eliminated by removing the two edge rows and columns from the analysis. SUDHL4 cells were seeded at 10,000 cells/well in 25µL of RPMI-1640 + 20% heat-inactivated FBS using the Biotek MultiFloFX Dispenser. Plates were sealed with Breathe-easy membrane (Sigma-Aldrich) and incubated at 37 °C and 5% CO_2_ for 48 hours. 0.2 µg/mL rituximab was added using Echo liquid handler and incubated for 30 minutes. 20% human serum (f.v.) was added using the Biotek MultiFloFX dispenser. After two hours of incubation, cell viability was measured using Cell Titer Glo (Promega) on a microplate reader (Tecan). The readout was normalized among duplicate replicates.

### Chromatin immunoprecipitation (CHIP)

Approximately 10 million cells were crosslinked with 1% formaldehyde (final concentration) for 10 min at room temperature, and the reaction was quenched with 250 mM glycine. All subsequent steps were performed on ice or at 4 °C. Cell pellets were lysed sequentially in Lysis Buffer 1 (LB1: 50 mM HEPES-KOH (pH 7.5), 140 mM NaCl, 1 mM EDTA (pH 8.0), 10% glycerol, 0.5% NP-40, 0.25% Triton X-100), Lysis Buffer 2 (LB2: 10 mM Tris-HCl (pH 8.0), 200 mM NaCl, 1 mM EDTA (pH 8.0), 0.5 mM EGTA (pH 8.0)), and Lysis Buffer 3 (LB3: 10 mM Tris-HCl, 1 mM EDTA (pH 8.0), 0.1% SDS buffer), all lysis buffers containing protease inhibitor cocktail. After each buffer incubation (10 minutes on ice), cells were pelleted and the supernatant discarded. Chromatin was sonicated using a Bioruptor Pico (Diagenode) at 30 seconds on/off cycles for 12 cycles, to shear DNA to approximately 150–300 bp fragments. Sonicated lysates were clarified by centrifugation at 20,000 × g for 15 minutes at 4 °C.

Antibody coupling was performed by an overnight incubation of Dynabeads Protein A (Thermo Fisher Scientific) with Sp1 [EPR22648;50] CHIP grade (Abcam, ab231778) at 4 °C with rotation. Antibody-conjugated beads were washed three times with LB3 and incubated with sheared chromatin overnight at 4 °C on a rotating platform. Beads were then washed sequentially with low-salt sonication buffer (twice), high-salt sonication buffer, LiCl buffer, and TE buffer. Each wash involved gentle mixing, incubation on ice for 3–5 minutes, and brief centrifugation followed by magnetic separation. Chromatin was eluted by resuspending beads in 100 µL elution buffer and incubating at 65 °C for 30 minutes with gentle vortexing every 10 minutes. Beads were pelleted and eluates transferred to fresh tubes. WCL aliquots were thawed and processed in parallel. Crosslinks were reversed by overnight incubation at 65 °C with NaCl and RNase A. The following day, samples were treated with Proteinase K and incubated at 55 °C for 45 minutes.

DNA was purified using the Zymo DNA Clean & Concentrator kit according to the manufacturer’s instructions and eluted in 12 µL nuclease-free water. Fragment size distribution was assessed by agarose gel electrophoresis (1% agarose, EtBr) or Agilent Bioanalyzer, with WCL diluted 1:10 prior to analysis.

### Immunoprecipitation

To isolate the nuclear fraction, cells were resuspended in buffer A (10mM HEPES-NaOH pH 7.9, 10mM KCl, 1.5mM MgCl2, 0.34M sucrose, 10% glycerol, 0.5mM DTT) plus protease inhibitor and phosphatase inhibitor tablets for 10 minutes on ice at 4°C and centrifuged (1300xg, 5 minutes). The nuclear pellets were then resuspended with Buffer B (3mM EDTA, 0.2mM EGTA, 1mM DTT, 0.3U/µL Benzonase) plus protease inhibitor and phosphatase inhibitor tablets for 30 minutes on ice at 4°C. 400µg of protein, determined by the BCA Protein Assay Kit (Thermo Scientific), was incubated with Sp1 [EPR22648;50] CHIP grade (Abcam, ab231778) or Rabbit Isotype IgG Control (Invitrogen, #02-6102) in IP Lysis buffer (10mM Tris pH 7.4, 150mM NaCl, 0.5mM EDTA pH 8, 0.5% NP-40, 0.25% sodium deoxycholate, 2mM MgCl2) plus protease inhibitor and phosphatase inhibitor tablets, overnight at 4°C on a rotating wheel. The lysate was then incubated with Dynabeads™ Protein A (Invitrogen) for 3 hours at 4°C on a rotating wheel, washed 4 times with IP buffer (10mM Tris pH 7.4, 150mM NaCl, 0.5mM EDTA pH 8, 0.5% NP-40, 0.25% sodium deoxycholate), and eluted with 2x Laemmli and beta-mercaptoethanol elution buffer by heating to 95°C for 5 minutes. Sp1 IP samples were analysed by label-free quantification mass spectrometry essentially as described in (47).

### Label-free mass spectrometry (LFQ-MS)

To this end, eluted proteins were separated by SDS-PAGE gel electrophoresis using NuPAGE 12 % Bis-Tris Mini Protein gels (Thermo Scientific) at 170 V for 10 minutes. Each sample was treated as a single fraction for in-gel trypsin digestion. Briefly, gel fractions were cut approximately into 2 by 2 by 1 mm pieces, destained twice using 1 ml Destaining Solution (50 % (v/v) ethanol, 25 mM ammonium bicarbonate) and dehydrated with 150 ul acetonitrile. Gel pieces were dried using a speed vacuum concentrator (Eppendorf) and subsequently treated with 200 ul Reducing Solution (10 mM DTT, 50 mM ammonium bicarbonate) at 56 °C for 1 hour followed by 200 ul Alkylating Solution (55 mM iodoacetamide, 50 mM ammonium bicarbonate) at room temperature for 45 minutes in the dark. Gel pieces were washed once with 200 ul 50 mM ammonium bicarbonate and twice with 150 ul acetonitrile, dried using a speed vacuum concentrator, and incubated with 2 μg sequencing grade trypsin (Promega) in 200 ul 50 mM ammonium bicarbonate at 37 °C overnight. Digested peptides were extracted from the gel pieces on the following day through two incubations each using an alternating cycle of 200 ul Extraction Buffer (3 % (v/v) trifluoroacetic acid, 30 % (v/v) acetonitrile, 33.3 % (v/v) mM ammonium bicarbonate) and 150 ul acetonitrile for 20 minutes and 10 minutes, respectively, followed by a final incubation in 150 ul acetonitrile for 10 minutes. All supernatants were collected and combined. Extracted peptides in solution were concentrated using a speed vacuum concentrator for 2 hours and desalted with home-made stage-tips containing C-18 resins (Empore). Stage tips were stored at 4°C prior elution of peptides with 30 ul Solution B (80 % (v/v) acetonitrile, 0.1% (v/v) formic acid) for analysis using an EASY-nLC 1200 Liquid Chromatograph (Thermo Scientific) coupled to a Q Exactive HF (Thermo Scientific) mass spectrometer.

Peptides were separated on a C-18-reversed phase column (25 cm length, 75 μm inner diameter; New Objective) which was packed in-house with ReproSil-Pur 120 C18-AQ 1.9 μm resin (Dr Maisch). The column was mounted on an Easy Flex Nano Source and maintained at 40 °C by a column oven (Sonation). An 88-minute gradient from 2 to 40 % Solution B at a flow of 225 nl/minute was used, followed by a 12 minute washout with 95 % Solution B. The spray voltage was set to 2.2 kV. The Q Exactive HF was operated with a TOP20 MS/MS spectra acquisition method per MS full scan, conducted with 60,000 at a maximum injected time of 20 ms and MS/MS scans with 15,000 resolution at a maximum injection time of 75 ms.

Raw files were processed with MaxQuant (48) version 2.0.1.0 with preset standard settings for label-free quantitation using at least 2 LFQ ratio counts based on the MaxLFQ algorithm (49). Carbamidomethylation was set as fixed modification while methionine oxidation, protein N-acetylation, and serine, threonine and lysine (STY) phosphorylation were considered as variable modifications. Searches were performed against the human (UP000005640) Uniprot databases. Search results were filtered with a false discovery rate of 0.01. Known contaminants, proteins groups only identified by site, and reverse hits of the MaxQuant results were removed, and only protein entries with at least three of four non-zero values in either condition of each pair-wise comparison were included in the analyses. Missing LFQ intensity values were imputed three times using a random distribution centered around the lowest 5% of LFQ intensities, with the final values representing the mean of these three iterations.

### Complement-dependent cytotoxicity assay

Cells were incubated with rituximab (RTX) for 30 minutes in a 96-well solid white flat bottom plate (Corning) at a cell density of 2*10^5^ cells/mL. SUDHL4 cells were treated with 37ng/mL RTX, while SUDHL6 cells were treated with 12.5 ng/mL RTX unless indicated. Subsequently, pooled human complement serum (cat: ICSER100ML; iDNA) or heat-inactivated (complement-inactivating) serum was added to 10% v/v concentration and incubated for 2 hours. Cell viability was determined by Cell Titer Glo (Promega) with SparkControl V2.3 Tecan Microplate Reader at 1s integration. Percent direct death was calculated as [(1-(RTX+hiHS / HS only) x 100)]. Percent CDC was calculated as [(1-(RTX+HS / HS only)) x 100) -% direct death]. “RTX” represents rituximab, “HS” represents human complement serum, and “HiHS” represents heat inactivated human complement serum.

For mCRP neutralizing experiments, CD55 antibody (BRI216) (Bio-Rad, MCA914) or CD59 (YTH53.1) (Bio-rad, MCA715G) were added at 10µg/mL final concentration for 1 hour prior to the complement assay.

### Antibody-dependent cellular cytotoxicity assay

Antibody-dependent cell-mediated cytotoxicity (ADCC) was assessed using a fluorescence imaging-based assay adapted from established flow cytometry protocols (50). Target cells were labeled with 2 μM carboxyfluorescein succinimidyl ester (CFSE) for 10 minutes at 37°C. Following washing, CFSE-labeled target cells (10,000 per well) were seeded in 96-well glass-bottom imaging plates and pre-incubated with 10ng/mL Rituximab for 30 minutes at 37°C. Effector NK-92.CD16 cells were then added at 1:1 effector-to-target ratio. To assess cell viability and cytotoxicity, 4’,6-diamidino-2-phenylindole (DAPI) was added to a final concentration of 0.2 μg/ml at a 4-hour timepoint, allowing 5 minutes for uptake before image acquisition. Cytotoxicity was quantified by counting CFSE-positive target cells that co-stained with DAPI, indicating membrane permeabilization and cell death. Percent cell death was calculated as: [(CFSE^+^ DAPI^+^ cell count / CFSE^+^ cell count) × 100]. Controls included target cells with Rituximab, and target and NK cells with isotype-matched control antibodies.

### Flow cytometry

Cells were stained with conjugated antibodies according to manufacturer’s instructions. Briefly, 1 × 10^6 cells were stained in a 100-µl experimental sample for 30 minutes on ice and washed twice prior to analysis. Flow cytometry was performed using the BD ® LSR II Flow Cytometer. Debris was excluded by gating out low FCS-A and SSC-A cells. Single cell populations were obtained by gating out FCS-H-low cells in a FCS-H v FSC-A plot. Cells were analyzed with FlowJo™ v10.8 Software (BD Life Sciences). Mean fluorescence intensity (MFI) was calculated using the median for unimodal cell populations. Integrated mean fluorescence intensity (iMFI) was calculated for bimodal populations (51). Relative MFI was calculated by normalizing each marker to the respective IgG control, then comparing the treatment to control condition (47). Antibodies specific for the following proteins were used for flow cytometry: APC mouse anti-human CD20 (BD Bioscience, #559776), APC mouse IgG1, κ Isotype control (BD Bioscience, #555751), PE mouse anti-human CD46 (BD Bioscience, #56424), PE mouse anti-human CD55 (BD Bioscience, #341030), PE mouse anti-human CD59 (BD Bioscience, #553457), PE mouse IgG2a, κ Isotype Control (BD Bioscience, #553457).

### RNA extraction and quantitative reverse transcription–PCR (qRTDPCR)

Total RNA was extracted using RNeasy Mini kit (Qiagen) according to manufacturer’s instructions and was reverse-transcribed to cDNA using iScript cDNA synthesis kit (BioRad). Quantitative real-time PCR was performed with PrecisionFAST qPCR Master Mix (Primerdesign Ltd) and primers (IDT, listed in Materials), and mRNA expression levels of the genes of interest were evaluated by the ΔΔ Ct method after qPCR analysis on the Comparative-CT program on QuantStudio 3 and 5 Real-Time PCR system (Fisher Scientific). Endogenous 18S was regarded as internal control. qPCR primers are detailed in Supplementary Materials.

Primers were designed to amplify a region of the *CD59* gene promoter containing putative SP1 transcription factor binding sites, based on genomic coordinates from the human reference genome (GRCh38/hg38). The forward and reverse primers were 5′-GTGCTCATTGGGTCCTGGCCAC-3′ and 5′-GCGCCCAAGATCCTCTTCCAGC-3′, respectively. This primer pair amplifies a 248 bp region of the *CD59* promoter (chr11:33,736,388–33,736,635; GRCh38). The amplicon was scanned for SP1 binding motifs using the JASPAR database (SP1 matrix family MA0079), and several high-confidence sites (relative scores ≥0.9) were identified: GGGGCGGGG (MA0079.5) at chr11:33,736,539– 33,736,547 (+ strand); GCTCCGCCCCC (MA0079.3) at chr11:33,736,551–33,736,561 (– strand); CCTCCACCCCC (MA0079.3) at chr11:33,736,452–33,736,462 (– strand). These motifs are located within a GC-rich region of the *CD59* promoter, consistent with known SP1 binding preferences, supporting the rationale for selecting this fragment for SP1 ChIP-qPCR analysis.

### Gene expression analyses across tumor tissue samples

From public datasets, we analyzed gene expression of complement regulatory proteins (*CD46*, *CD55*, and *CD59*) in DLBCL cell lines (DB, Karpas422, RCK8, SUDHL10, U2932, WSUDLCL2, OCILY1, OCILY18, RI1, SUDHL2, SUDHL4, TOLEDO, CARNAVAL, DOHH2, NUDHL1, OCILY7, SUDHL5, SUDHL6) following doxorubicin treatment, and investigated *CD59* expression in patient tissue samples before and after chemotherapy. Data were obtained from the GEO database and analyzed using GEO2R, except for GSE191127 and GSE224867, which underwent independent processing. The following datasets were analyzed: GSE224867 (RNA-seq of DLBCL cell lines treated with/without doxorubicin; Illumina TruSeq Stranded RNA libraries sequenced at DKFZ, processed using FastQC v0.11.9, Salmon v1.5.2, and DESeq2 v1.34.0); GSE15781 (42 colorectal cancer samples with pre-operative neoadjuvant radiochemotherapy (RCT), including 9 irradiated tumors, 13 non-irradiated tumors, 10 irradiated normal tissues, and 10 non-irradiated normal tissues; log2-transformed GPL2986 microarray data); GSE227100 (RNA-seq of 24 paired pre-and post-treatment HGSOC metastatic tumor samples; Illumina NovaSeq 6000, Kallisto quantification (TPM)); GSE146963 (microarray data of 28 paired pre- and post-treatment HGSOC tumor samples; SST-RMA normalized Affymetrix data); GSE191127 (RNA-seq of 20 paired pre- and post-treatment HER2-negative breast cancer biopsies; Illumina HiSeq 2000, DESeq2 normalized and log2-transformed); and GSE21974 (32 paired primary invasive breast cancer core biopsies before and after 4 cycles of neoadjuvant chemotherapy with epirubicine and cyclophosphamide followed by docetaxel; log2-transformed Agilent-014850 44K single-color microarray data processed using Agilent Feature Extraction Software v9.1). Statistical comparisons were performed using Wilcoxon tests for paired samples and unpaired t-tests for unpaired samples.

## Statistical analysis

Statistical analyses and graphing were performed using GraphPad Prism 10.1.1 software (Graphpad, San Diego, CA, USA). For comparisons of two groups, statistical analysis was performed using paired t-tests, while ANOVA followed by post hoc Dunnet’s multiple comparisons test to etoposide condition was used for multiple groups. Error bars represent SD of biological replicates. Statistical significance was denoted as follows (*) p < 0.05, (**) p < 0.01, (***) p < 0.001.

## Supplementary Materials and Figures

Materials

Fig. S1 Phage-display Antibody Screen Design

Fig. S2 Etoposide-induced CD59 gene expression is rescued with Mithramycin A

Table S1 Co-immunoprecipitated proteins pulled down with SP1 vs IgG antibodies, identified by mass spectrometry within CTRL condition.

Table S2 Co-immunoprecipitated proteins pulled down with SP1 vs IgG antibodies, identified by mass spectrometry within ETO condition.

Table S3 Co-immunoprecipitated proteins pulled down with SP1 vs IgG antibodies, identified by mass spectrometry within ETO+Chk1i condition.

## Acknowledgments

Donut charts and heatmaps were generated using Rstudio (V4.4.0). Visuals were created with Biorender.com.

## Acknowledgements

We thank Michelle Mok Meng Huang and Erica Xu Songci from the Fluorescence Activated Cell Sorting Facility at CSI for maintenance of flow cytometry machines and guidance on flow cytometry analysis. We thank Choong Meng Ling from the Experimental Drug Development Centre at A*STAR for assistance in the design of the chemical screen. We thank Giridharan Periyasamy, Matan Thangavelu, and Shivaji Rikka from Center for High-throughput Phenomics, Genome Institute of Singapore for providing access to the high-throughput screening facility and drug libraries, and discussion on design of chemical screen. We thank Dr. Joseph J Zhao from the Yong Loo Lin School of Medicine, National University of Singapore for assistance with Rstudio Figs.

## Author contributions

Conceptualization: ASYC, ADJ

Methodology: ASYC, WKY, VK, LJM, NM, DK, ADJ

Validation: ASYC, AA, CZYO, MMH, NM, MIBA, PMH, GH

Formal analysis: ASYC, VK

Investigation: ASYC, AA, CZYO, MMH, NM, GH, ADJ

Writing – Original Draft: ASYC, ADJ

Writing – Review & Editing: ASYC, PWJ, WKY, NM, WJC, MSC, DK, CT, ADJ

Visualization: ASYC, VK

Supervision: ADJ

Project administration: PWJ, ADJ

Funding acquisition: CT, ADJ

## Data and materials availability

All data relevant to the study are included in the article or uploaded as online supplemental information. The mass spectrometry data have been deposited to the ProteomeXchange Consortium via the PRIDE partner repository with the dataset identifier PXD056654 (52).

**Figure S1.**
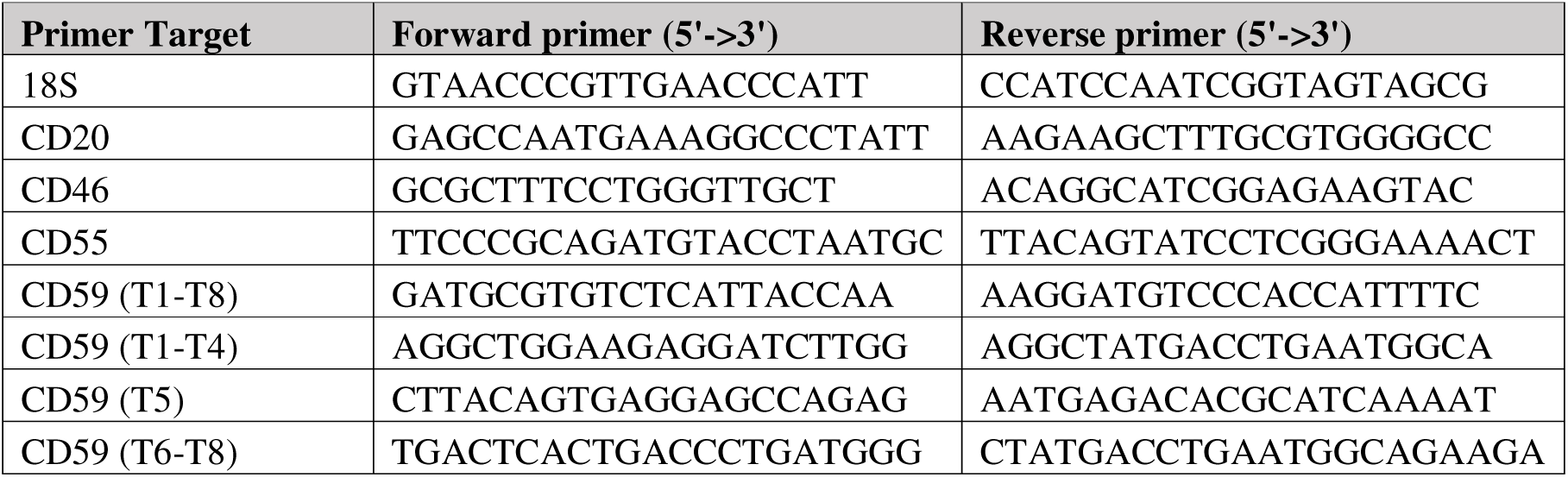
Phage-display Antibody Screen Design. Schematic illustration of the multi-stage process of the phage-display antibody screen used to identify antibodies that preferentially bind to DNA-damaged cells. **(A)** The process begins with the construction of the phage-display antibody library, which includes generating a diverse library of antibody sequences, cloning these sequences, and producing the antibody phage library. **(B)** In the initial screening step, cells exposed to the phage library were treated with an organic solvent and sedimented to remove non-specific binders. The phage library was then exposed to undamaged MCF10A cells, and the binders obtained through ICOS (Immunoprecipitation of Crosslinked and Sonicated) were converted into a masking antibody set for MCF10A cells. **(C)** The final stage involves the screening of DNA damage-specific binders. In this step, irradiated MCF10A cells were treated with the masking antibody set and then exposed to the phage library to obtain phage complexes by ICOS that preferentially bind DNA-damaged cells. To validate the results, scFv fragments were purified from 20 selected phage complexes and tested by flow cytometry to select the best candidates based on consistency of DNA damage-related induction. This phage-display screen design allows for the identification of antibodies that specifically recognize epitopes present or upregulated in DNA-damaged cells while minimizing binding to undamaged cells.

**Figure S2.** Etoposide-induced *CD59* gene expression is rescued with Mithramycin A. **(A)** Schematic diagram of CD59 gene transcriptional regulators (adapted from Du et al.). Sp1 and NF-κB primarily regulate transcripts T1-T4. TP53 regulates the T5 transcript but has minimal impact on overall CD59 expression. CREB primarily regulates T6-T8 transcripts and plays additional roles in CD59 transcription beyond T6-T8 regulation. An enhancer region (- 500 to -1000 bp) interacts with multiple factors (CREB, NF-κB, TP53) through CBP/p300 scaffolding, facilitating coordinated regulation of CD59 expression. **(B)** Relative mRNA expression of *CD59* exons T1-8, T1-4, T5, and T6-8 in SUDHL6 cells after ETO or ETO + Chk1i treatment. **(C)** Percentage of CDC in SUDHL4 cells treated with indicated NF-κB inhibitors for 48 hours. (n=3 technical triplicates). **(D)** Percentage of CDC in SUDHL6 cells pretreated with etoposide (ETO) alone or in combination with Mithramycin A (MitA) (n=3 biological replicates). **(E)** Relative CD46 expression in SUDHL4 cells treated with etoposide alone or in combination with Mithramycin A (n=2 biological replicates). **(F)** Flow cytometry histogram of SUDHL4 cells treated with or without Mithramycin A, showing IgG-PE control and CD46-PE staining. Statistical analysis for all panels was performed using one-way ANOVA followed by Dunnett’s multiple comparisons test, with the etoposide-treated condition as the reference group. Error bars represent standard deviation. Statistical significance is denoted as follows: * p < 0.05, ** p < 0.01, *** p < 0.001.

## References

1. Klapp V, Álvarez-Abril B, Leuzzi G, Kroemer G, Ciccia A, Galluzzi L. The DNA damage response and inflammation in cancer. Cancer Discov. American Association for Cancer Research (AACR); 2023;13:1521–45.

2. Liu B, Zhou H, Tan L, Siu KTH, Guan X-Y. Exploring treatment options in cancer: Tumor treatment strategies. Signal Transduct Target Ther. Springer Science and Business Media LLC; 2024;9:175.

3. Sato H, Niimi A, Yasuhara T, Permata TBM, Hagiwara Y, Isono M, et al. DNA double-strand break repair pathway regulates PD-L1 expression in cancer cells. Nat Commun. 2017;8:1751.

4. Uchihara Y, Permata TBM, Sato H, Kawabata-Iwakawa R, Katada S, Gu W, et al. DNA damage promotes HLA class I presentation by stimulating a pioneer round of translation-associated antigen production. Mol Cell. Cell Press; 2022;82:2557–2570.e7.

5. Gasser S, Orsulic S, Brown EJ, Raulet DH. The DNA damage pathway regulates innate immune system ligands of the NKG2D receptor. Nature. 2005;436:1186–90.

6. Kurosawa M, Jeyasekharan AD, Surmann E-M, Hashimoto N, Venkatraman V, Kurosawa G, et al. Expression of LY6D is induced at the surface of MCF10A cells by X-ray irradiation. FEBS J. 2012;279:4479–91.

7. Akahori Y, Kurosawa G, Sumitomo M, Morita M, Muramatsu C, Eguchi K, et al. Isolation of antigen/antibody complexes through organic solvent (ICOS) method. Biochem Biophys Res Commun. Elsevier BV; 2009;378:832–5.

8. Walport MJ. Complement. First of two parts. N Engl J Med. 2001;344:1058–66.

9. Kojima A, Iwata K, Seya T, Matsumoto M, Ariga H, Atkinson JP, et al. Membrane cofactor protein (CD46) protects cells predominantly from alternative complement pathway-mediated C3-fragment deposition and cytolysis. J Immunol. The American Association of Immunologists; 1993;151:1519–27.

10. Medof ME, Kinoshita T, Nussenzweig V. Inhibition of complement activation on the surface of cells after incorporation of decay-accelerating factor (DAF) into their membranes. J Exp Med. Rockefeller University Press; 1984;160:1558–78.

11. Meri S, Morgan BP, Davies A, Daniels RH, Olavesen MG, Waldmann H, et al. Human protectin (CD59), an 18,000-20,000 MW complement lysis restricting factor, inhibits C5b-8 catalysed insertion of C9 into lipid bilayers. Immunology. 1990;71:1–9.

12. Roumenina LT, Daugan MV, Petitprez F, Sautès-Fridman C, Fridman WH. Context-dependent roles of complement in cancer. Nat Rev Cancer. Springer Science and Business Media LLC; 2019;19:698–715.

13. Yamamoto H, Fara AF, Dasgupta P, Kemper C. CD46: the “multitasker” of complement proteins. Int J Biochem Cell Biol. Elsevier BV; 2013;45:2808–20.

14. Bharti R, Dey G, Lin F, Lathia J, Reizes O. CD55 in cancer: Complementing functions in a non-canonical manner. Cancer Lett. Elsevier BV; 2022;551:215935.

15. Thi TN, Thanh HD, Nguyen V-T, Kwon S-Y, Moon C, Hwang EC, et al. Complement regulatory protein CD46 promotes bladder cancer metastasis through activation of MMP9. Int J Oncol [Internet]. Spandidos Publications; 2024;65. Available from: 10.3892/ijo.2024.5659

16. Dho SH, Lim JC, Kim LK. Beyond the role of CD55 as a complement component. Immune Netw. 2018;18:e11.

17. Geller A, Yan J. The Role of Membrane Bound Complement Regulatory Proteins in Tumor Development and Cancer Immunotherapy. Front Immunol. Frontiers; 2019;10:440775.

18. Weiner GJ. Rituximab: mechanism of action. Semin Hematol. Elsevier BV; 2010;47:115–23.

19. Cragg MS, Morgan SM, Chan HTC, Morgan BP, Filatov AV, Johnson PWM, et al. Complement-mediated lysis by anti-CD20 mAb correlates with segregation into lipid rafts. Blood. 2003;101:1045–52.

20. Golay J, Taylor RP. The Role of Complement in the Mechanism of Action of Therapeutic Anti-Cancer mAbs. Antibodies (Basel) [Internet]. 2020;9. Available from: 10.3390/antib9040058

21. Hu W, Ge X, You T, Xu T, Zhang J, Wu G, et al. Human CD59 inhibitor sensitizes rituximab-resistant lymphoma cells to complement-mediated cytolysis. Cancer Res. American Association for Cancer Research (AACR); 2011;71:2298–307.

22. Du Y, Teng X, Wang N, Zhang X, Chen J, Ding P, et al. NF-κB and enhancer-binding CREB protein scaffolded by CREB-binding protein (CBP)/p300 proteins regulate CD59 protein expression to protect cells from complement attack. J Biol Chem. 2014;289:2711–24.

23. Kurosawa G, Akahori Y, Morita M, Sumitomo M, Sato N, Muramatsu C, et al. Comprehensive screening for antigens overexpressed on carcinomas via isolation of human mAbs that may be therapeutic. Proc Natl Acad Sci U S A. 2008;105:7287–92.

24. Candelaria M, Dueñas-Gonzalez A. Rituximab in combination with cyclophosphamide, doxorubicin, vincristine, and prednisone (R-CHOP) in diffuse large B-cell lymphoma. Ther Adv Hematol. SAGE Publications; 2021;12:2040620721989579.

25. Belenki D, Richter-Pechanska P, Shao Z, Bhattacharya A, Lau A, Nabuco Leva Ferreira de Freitas JA, et al. Senescence-associated lineage-aberrant plasticity evokes T-cell-mediated tumor control. Nat Commun. Springer Science and Business Media LLC; 2025;16:3079.

26. van Meerten T, van Rijn RS, Hol S, Hagenbeek A, Ebeling SB. Complement-induced cell death by rituximab depends on CD20 expression level and acts complementary to antibody-dependent cellular cytotoxicity. Clin Cancer Res. 2006;12:4027–35.

27. Macor P, Tripodo C, Zorzet S, Piovan E, Bossi F, Marzari R, et al. In vivo targeting of human neutralizing antibodies against CD55 and CD59 to lymphoma cells increases the antitumor activity of rituximab. Cancer Res. American Association for Cancer Research (AACR); 2007;67:10556–63.

28. Snipstad K, Fenton CG, Kjaeve J, Cui G, Anderssen E, Paulssen RH. New specific molecular targets for radio-chemotherapy of rectal cancer. Mol Oncol. Wiley; 2010;4:52–64.

29. Adzibolosu N, Alvero AB, Ali-Fehmi R, Gogoi R, Corey L, Tedja R, et al. Immunological modifications following chemotherapy are associated with delayed recurrence of ovarian cancer. Front Immunol. 2023;14:1204148.

30. Jiménez-Sánchez A, Cybulska P, Mager KL, Koplev S, Cast O, Couturier D-L, et al. Unraveling tumor-immune heterogeneity in advanced ovarian cancer uncovers immunogenic effect of chemotherapy. Nat Genet. Springer Science and Business Media LLC; 2020;52:582–93.

31. Hoogstraat M, Lips EH, Mayayo-Peralta I, Mulder L, Kristel P, van der Heijden I, et al. Comprehensive characterization of pre-and post-treatment samples of breast cancer reveal potential mechanisms of chemotherapy resistance. NPJ Breast Cancer. Springer Science and Business Media LLC; 2022;8:60.

32. Stickeler E, Pils D, Klar M, Orlowsk-Volk M, Zur Hausen A, Jäger M, et al. Basal-like molecular subtype and HER4 up-regulation and response to neoadjuvant chemotherapy in breast cancer. Oncol Rep. Spandidos Publications; 2011;26:1037–45.

33. Martin BK. Transcriptional control of complement receptor gene expression. Immunol Res. Springer Science and Business Media LLC; 2007;39:146–59.

34. Torabi B, Flashner S, Beishline K, Sowash A, Donovan K, Bassett G, et al. Caspase cleavage of transcription factor Sp1 enhances apoptosis. Apoptosis. 2018;23:65–78.

35. Arede L, Pina C. Buffering noise: KAT2A modular contributions to stabilization of transcription and cell identity in cancer and development. Exp Hematol. Elsevier BV; 2021;93:25–37.

36. Gabriel GC, Yagi H, Tan T, Bais A, Glennon BJ, Stapleton MC, et al. Mitotic block and epigenetic repression underlie neurodevelopmental defects and neurobehavioral deficits in congenital heart disease. Nat Commun. Springer Science and Business Media LLC; 2025;16:469.

37. Golay J, Zaffaroni L, Vaccari T, Lazzari M, Borleri G-M, Bernasconi S, et al. Biologic response of B lymphoma cells to anti-CD20 monoclonal antibody rituximab in vitro: CD55 and CD59 regulate complement-mediated cell lysis. Blood. American Society of Hematology; 2000;95:3900–8.

38. Jiang K, Deng M, Du W, Liu T, Li J, Zhou Y. Functions and inhibitors of CHK1 in cancer therapy. Med Drug Discov. Elsevier BV; 2024;22:100185.

39. Kristeleit R, Plummer R, Jones R, Carter L, Blagden S, Sarker D, et al. A Phase 1/2 trial of SRA737 (a Chk1 inhibitor) administered orally in patients with advanced cancer. Br J Cancer. 2023;129:38–45.

40. Jones R, Plummer R, Moreno V, Carter L, Roda D, Garralda E, et al. A Phase I/II trial of oral SRA737 (a Chk1 inhibitor) given in combination with low-dose gemcitabine in patients with advanced cancer. Clin Cancer Res. American Association for Cancer Research (AACR); 2023;29:331–40.

41. Fan Z, Luo H, Zhou J, Wang F, Zhang W, Wang J, et al. Checkpoint kinase[1 inhibition and etoposide exhibit a strong synergistic anticancer effect on chronic myeloid leukemia cell line K562 by impairing homologous recombination DNA damage repair. Oncol Rep. Spandidos Publications; 2020;44:2152–64.

42. Isono M, Okubo K, Asano T, Sato A. Inhibition of checkpoint kinase 1 potentiates anticancer activity of gemcitabine in bladder cancer cells. Sci Rep. Springer Science and Business Media LLC; 2021;11:10181.

43. Trabolsi A, Arumov A, Schatz JH. Bispecific antibodies and CAR-T cells: dueling immunotherapies for large B-cell lymphomas. Blood Cancer J. Springer Science and Business Media LLC; 2024;14:27.

44. Lu Y, Zhao Q, Liao J-Y, Song E, Xia Q, Pan J, et al. Complement signals determine opposite effects of B cells in chemotherapy-induced immunity. Cell. Elsevier BV; 2020;180:1081–1097.e24.

45. Stadlbauer K, Andorfer P, Stadlmayr G, Rüker F, Wozniak-Knopp G. Bispecific mAb2 antibodies targeting CD59 enhance the complement-dependent cytotoxicity mediated by rituximab. Int J Mol Sci. MDPI AG; 2022;23:5208.

46. Chai SC, Goktug AN, Chen T. Assay validation in high throughput screening – from concept to application. Drug Discovery and Development - From Molecules to Medicine. InTech; 2015.

47. Manso BA, Medina KL. Standardized flow-cytometry-based protocol to simultaneously measure transcription factor levels. STAR Protoc. 2021;2:100485.

48. Cox J, Mann M. MaxQuant enables high peptide identification rates, individualized p.p.b.-range mass accuracies and proteome-wide protein quantification. Nat Biotechnol. Springer Science and Business Media LLC; 2008;26:1367–72.

49. Cox J, Hein MY, Luber CA, Paron I, Nagaraj N, Mann M. Accurate proteome-wide label-free quantification by delayed normalization and maximal peptide ratio extraction, termed MaxLFQ. Mol Cell Proteomics. Elsevier BV; 2014;13:2513–26.

50. Patra-Kneuer M, Chang G, Xu W, Augsberger C, Grau M, Zapukhlyak M, et al. Activity of tafasitamab in combination with rituximab in subtypes of aggressive lymphoma. Front Immunol. 2023;14:1220558.

51. Darrah PA, Patel DT, De Luca PM, Lindsay RWB, Davey DF, Flynn BJ, et al. Multifunctional TH1 cells define a correlate of vaccine-mediated protection against Leishmania major. Nat Med. 2007;13:843–50.

52. Perez-Riverol Y, Bai J, Bandla C, García-Seisdedos D, Hewapathirana S, Kamatchinathan S, et al. The PRIDE database resources in 2022: a hub for mass spectrometry-based proteomics evidences. Nucleic Acids Res. Oxford University Press (OUP); 2022;50:D543–52.

