## Supplementary figures and images for "Genotoxic chemotherapy impedes complement dependent cytotoxicity via Chk1-mediated CD59 regulation"

### Supplementary Figure 1

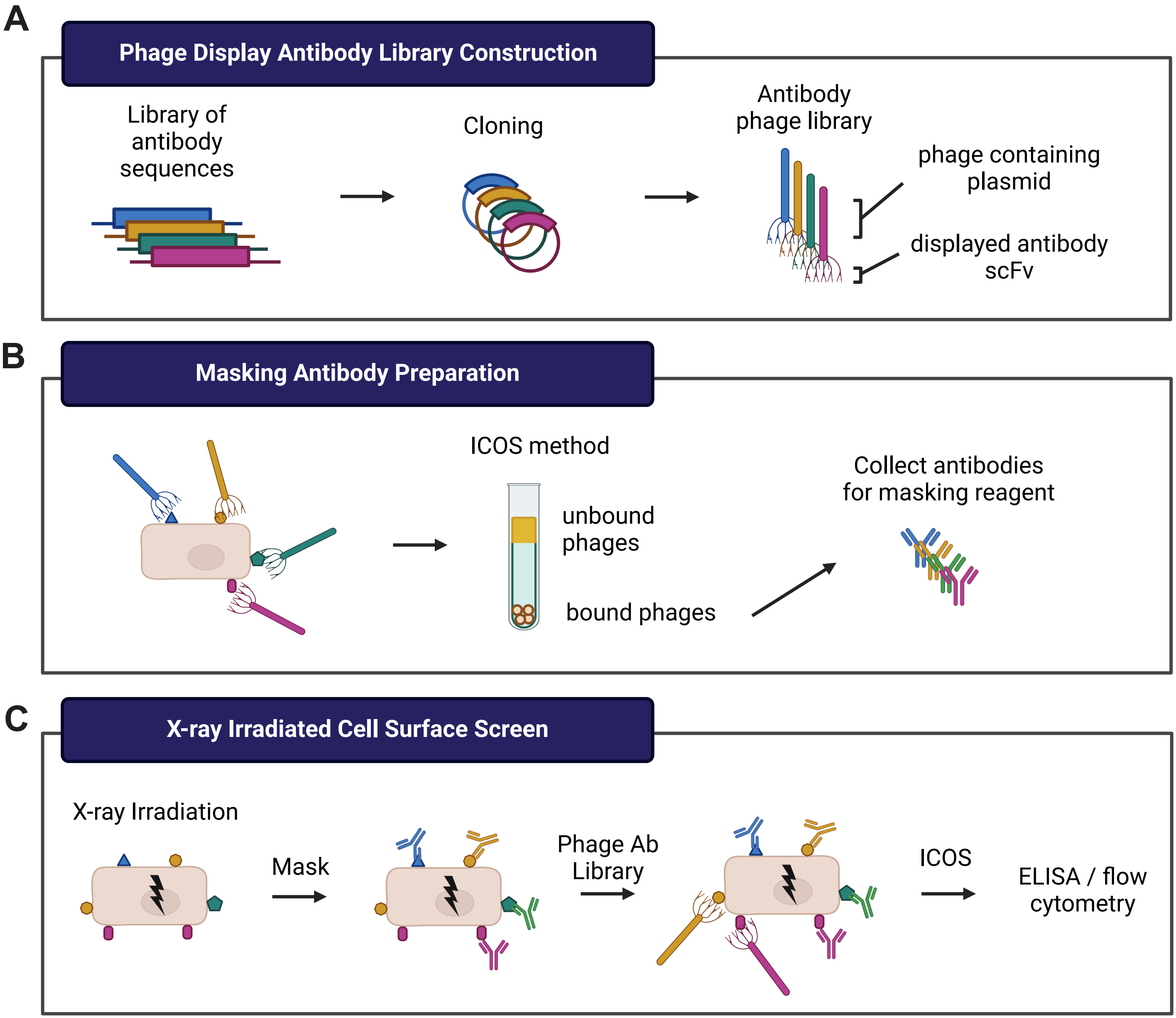

### Supplementary Figure 2

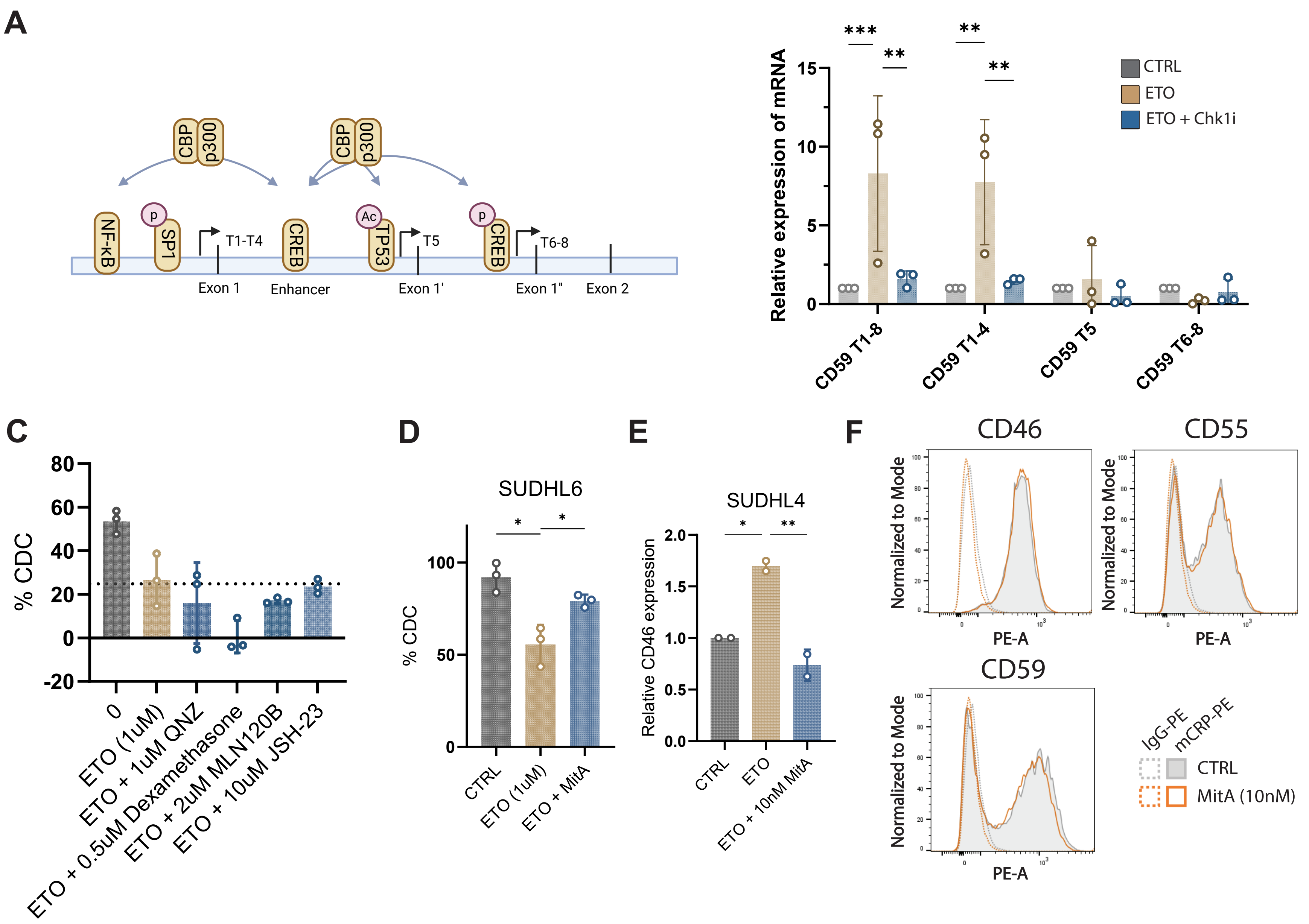
